# p73 is required for vessel integrity controlling endothelial junctional dynamics through Angiomotin

**DOI:** 10.1101/2022.03.03.482832

**Authors:** Laura Maeso-Alonso, Hugo Alonso-Olivares, Nicole Martínez-García, Lorena Lopez-Ferreras, Javier Villoch-Fernández, Laura Puente-Santamaría, Natalia Colas-Algora, Alfonso Fernández-Corona, María Elena Lorenzo-Marcos, Benilde Jiménez, Lars Holmgren, Margareta Wilhelm, Jaime Millan, Luis del Peso, Lena Claesson-Welsh, Margarita M. Marques, Maria C. Marin

## Abstract

Preservation of blood vessels integrity, which is critical for normal physiology and organ function, is controlled at multiple levels, including endothelial junctions. However, the mechanism that controls the adequate assembly of endothelial cell junctions is not fully defined. Here we uncover TAp73 transcription factor as a vascular architect that orchestrates transcriptional programs involved in cell junction establishment and developmental blood vessel morphogenesis and identify Angiomotin (AMOT) as a TAp73 direct transcriptional target. Knockdown of p73 in endothelial cells not only results in decreased Angiomotin expression and localization at intercellular junctions, but also affects its downstream function regarding Yes-Associated Protein (YAP) cytoplasmic sequestration upon cell-cell contact. Analysis of adherens junctional morphology after p73-knockdown in human endothelial cells revealed striking alterations, particularly a sharp increase in serrated junctions and actin bundles appearing as stress fibers, both features associated with enhanced barrier permeability. In turn, stabilization of Angiomotin levels rescued those junctional defects, confirming that TAp73 controls endothelial junction dynamics, at least in part, through the regulation of Angiomotin. The observed defects in monolayer integrity were linked to hyperpermeability and reduced transendothelial electric resistance. Moreover, p73-knockout retinas showed a defective sprout morphology coupled to hemorrhages, highlighting the physiological relevance of p73 regulation in the maintenance of vessel integrity *in vivo*. We propose a new model in which TAp73 acts as a vascular architect integrating transcriptional programs that will impinge with Angiomotin/YAP signaling to maintain junctional dynamics and integrity, whilst balancing endothelial cell rearrangements in angiogenic vessels.

## Introduction

Formation of blood vessel networks by sprouting angiogenesis is critical for tissue growth, homeostasis and regeneration. During this process, the rearrangements of tip- and stalk-cells require dynamic remodeling of cell-cell junctions while maintaining the integrity of the new sprout to ensure proper barrier functionality [1]. In this context, endothelial cells (EC) junctions play a coordinating role, polarizing the activity of the migration machinery and maintaining the sprout cohesion [2]. Loss of these junctions promotes hemorrhages and edemas and also plays an important role during tumor angiogenesis [3]. Junction-associated proteins, like the angiomotin family member Angiomotin (AMOT) are involved in the regulation and coordination of front–rear polarity and contribute to the assembly of EC junctions and the stabilization and maturation of vessels [4-7]; however, the identification of upstream regulators of these proteins remain elusive.

In this regard, the *TP73* gene has been described as an essential player during angiogenesis [8-10]. *TP73* can give rise to different functional isoforms: TAp73 with full transactivation activity, and the truncated dominant-negative DNp73 [11]. DNp73 functions as a pro-angiogenic signaling factor in tumor cells [9, 10, 12], whereas a bi- functional role in tumor angiogenesis has been proposed for TAp73 [8, 10, 13]. However, the mechanism of TAp73 physiological function during developmental vascular morphogenesis has not been unraveled. The *Trp73-/-* mice (p73KO), which lack both isoforms, exhibit extensive gastrointestinal and cranial hemorrhages [14], suggestive of vascular fragility and/or other defects in their vascular compartment. p73 deficiency also leads to perturbed vascular development in the mouse retina, with a decrease in vascular branching and elongation, and anastomosis defects [9]. Interestingly, similar defects have previously been associated with altered stability or dynamics of EC junctions [15-19]. However, whether TAp73-deficiency in EC leads to junctional defects with important consequences for vascular morphogenesis remains an important open question.

In this work we demonstrate p73 essential role in the maintenance of the endothelial junction dynamics and barrier integrity. Our results demonstrate that p73 regulates endothelial junction dynamics, at least in part, via AMOT, linking for the first time p73 and the AMOT/YAP pathway in the regulation of endothelial junctions. This situates p73 function in the hub of upstream regulators that control how endothelial cells establish and maintain adequate junctions to shape functional vascular networks with important consequences in the maintenance of vascular homeostasis.

## Material and methods

### Mice husbandry, animal breeding and genotyping

*Trp73*^-/-^ mice (p73KO) and DNp73 mutant mice (DNp73KO) were obtained as described before [20]. Male and female littermates were used for the experiments and genotyping was performed according to [14, 20].

### Retina dissection and flat-mount preparation

Retinal extraction and flat-mount preparation were performed as described in [9]. Eyes from postnatal mice on day 7 (P7) were extracted from either perfused or non-perfused mice. In the latter case, eyes were fixed with 4% PFA during 40 min and stained immediately. Wholemount (WM) staining was performed according to [9], using VE-cadherin (BD #555298, Clone 11D4.1, 1:100 dilution, retinas from perfused mice) or TER119 (R&Dsystems #MAB1125-SP, 1:100 dilution, retinas from non-perfused mice) primary antibodies, and a donkey anti-Rat Alexa Fluor 488 (Invitrogen, # A21208) as secondary antibody, as well as biotin-conjugated isolectin B4 (IB4, Sigma-Aldrich, #L2140).

### Cell culture

Generation and culture of murine wild type (WT) and p73KO-induced pluripotent stem cells (iPSCs) has been described elsewhere [21]. iPSCs were cultured on MEFs feeder cells (5 × 10^4^ cells/cm^2^) in Dulbecco’s Modified Eagle Medium (DMEM, Sigma-Aldrich, #D5671) supplemented with 15% FBS, 2 mM L-glutamine, 1 mM sodium pyruvate, 1 mM nonessential amino acids, 0.1 mM β-mercaptoethanol and 1000 U/ml Leukemia Inhibitory Factor (LIF, Millipore, #ESG1107).

Wild type E14TG2α mouse embryonic stem cells (E14-WT, mESCs) were kindly provided by Dr. Jim McWhir (former researcher at Roslin Institute, Edinburgh, Scotland, UK). TAp73-deficient cells (E14-TAp73KO) were generated by gene editing and cultured as previously described [22].

Human Umbilical Vein Endothelial Cells (HUVEC) were isolated from the veins of umbilical cords following the established protocol [23]. Cells were routinely maintained on 0.1% gelatine-coated flasks in endothelial basal medium, with full supplements (EBM Lonza, #CC3124). Cells were used for experiments between passage 4 to 8.

The human cell line HA-TAp73β-Saos2-Tet-On [24] was kindly provided by Dr. Karen Vousden (Cancer Research UK Beatson Institute, Glasgow), and cultured as described in [25]. For TAp73 induction, cells were treated with 2.5 μg/ml of doxycycline (DOXO) for 36 h.

### Three-dimensional (3D) endothelial differentiation

Endothelial differentiation assays were carried out as previously reported [9, 26]. Briefly, embryoid bodies (EBs) were formed by the hanging drop method and embedded in a collagen I matrix composed of Ham’s F12 medium (Sigma-Aldrich, #D8900), 6.26 mM NaOH, 12.5 mM HEPES, 0.073% NaHCO3, 1% GlutaMAX™ and 1.5 mg/ml collagen I (Advanced Biomatrix, #5005), supplemented with 100 ng/ml VEGF-A (Peprotech, #450-32). Sprouts were analysed after 12 days of *in vitro* culture (iPSCs) or at day 21 (mESCs).

### HUVEC Knockdown studies

Two different approaches were conducted to knockdown p73 expression. For synthetic small interfering RNA (siRNA), 5 × 10^5^ cells were nucleofected (Amaxa^®^ HUVEC Nucleofector^®^ Program A-034, Kit #VAPB-1002, Lonza) with 50 nM of the following siRNA oligos: scrambled (5′UAGCCACCACUGACGACCUdTdT3′, 5′dTdTAUCGGUGGUGACUGCUGGA3′), or sip73 (i4: 5′CCAUCCUGUACAACUUCAUGUG3′, 5′CAUGAAGUUGUACAGGAUGGUG3′) [27]. After nucleofection, cells were immediately seeded on 0.1% gelatine-coated 6-well plates and analysed after 48-96 h. Lentiviral vector-based RNA interference was carried out with a combination of two lentiviral vectors encoding short hairpin RNA to enhance the silencing efficiency: LV pGIPZ shp73ex5 (clone id V2LHS_69894) and LV pGIPZ shp73ex9 (clone id V3LHA_330455). An empty pGIPZ vector (Dharmacon, # RHS4349) was used as control (VECTOR). Lentiviruses at a multiplicity of infection (MOI) of 20 (10 MOI of each plasmid in case of shp73) were added to the culture medium (plus 8 μg/ml polybrene) for 8 h, resulting in more than 95% transduced cells (pGIPZ includes a GFP reporter). Seventy-two hours later, cells were seeded either at 7,5 × 10^4^ cells/cm^2^ (high density) or 1 × 10^4^ cells/cm^2^ (low density conditions) on 0.1% gelatine-coated plates and were analysed after 72 or 96 h.

For AMOT stabilization experiments, transduced cells were seeded at 7,5 × 10^4^ cells/cm^2^ and after 48-72 h, cells were treated with the tankyrase inhibitor XAV939 (5µM, Sigma-Aldrich, #CX3004) for 24 h.

### HUVEC Endothelial barrier function assays

The *in vitro* vascular permeability assay (Millipore, # ECM644-24 wells) was performed following the manufacturer’s instructions. Trans-endothelial electric resistance (TEER) assays with an electric cell-substrate impedance sensing system (ECIS 1600R; Applied Biophysics) [28] were performed as described [29]. Briefly, 150,000 infected cells were plated onto 8W10E PET arrays (ibidi #72001) and multifrequency measurement was carried out (ranges between 250 and 64000 Hz) for 48 h.

### Immunofluorescence and confocal microscopy image acquisition

Cells were fixed with 3.7% PFA (or 3.7% PFA + 0.5% Tx-100 in the case of ZO-1 staining) during 15 min at RT. After washing (1X PBS), cells were incubated for 15 min in permeabilization buffer (0.5% Tx100 in 1X PBS), blocked for 1 h (10% goat serum in PBS) and incubated with the corresponding primary antibodies at 4 ºC overnight (o/n) (Table 1). Then, cells were washed (1X PBS) and incubated for 45 min at RT with the secondary antibodies (Table 1). Finally, they were stained with DAPI (1 µg/ml), Phalloidin-TRICT (Sigma Aldrich P1951; 1:100 dilution) or Phalloidin-488 (Thermo Fisher Scientific A12379, 1:100) as indicated, and mounted with Fluoromount-G™ (Electron Microscopy Sciences #17984-25). The immunostaining of 3D-endothelial differentiated cells was performed following the protocol of (Fernandez-Alonso et al., 2015). Confocal microscopy images (8bits) were obtained in a Zeiss LSM800 Confocal Laser Scanning Microscope using 25x Plan-Apo/0.8 numerical aperture or 63x Plan-Apo/1.4 numerical aperture oil objectives at RT. In some cases, a 0.5 zoom was used. Confocal Z-stack images were acquired in all cases and stacks were z-projected to maximum intensity. Images were processed with the ZEN blue software (Carl Zeiss Microscopy GmbH).

**Table 1.** Primary and secondary antibodies used for immunofluorescence.

|  | Antibody | Host | Dilution | Manufacturer | Catalog Number |
| --- | --- | --- | --- | --- | --- |
| Primary | **VE-cadherin** | Goat | 1/100 | R&D Systems | AF1002 |
|  | **VE-cadherin (F8)** | Mouse | 1/500 | Santa Cruz Biotechnology | sc9989 |
|  | **YAP (63.7)** | Mouse | 1/100 | Santa Cruz Biotechnology | 101199 |
|  | **ZO-1 (1A12)** | Mouse | 1/100 | Thermo Fisher | 339100 |
|  | **β-Catenin (14)** | Mouse | 1/200 | BD transduction lab. | 61053 |
|  | **Amot (D2O4H)** | Rabbit | 1/200 | Cell Signalling | 43130S |
|  | **KI67** | Rabbit | 1/200 | Abcam | ab15580 |
| Secondary | **Alexa Fluor 488**<br>anti Rabbit | Donkey | 1/1000 | Molecular Probes | A21206 |
|  | **Alexa Fluor 488**<br>anti Mouse | Goat | 1/1000 | Molecular Probes | A21131 |
|  | **Alexa Fluor 546**<br>anti Mouse | Donkey | 1/1000 | Molecular Probes | A10036 |
|  | **Alexa Fluor IgG1 594**<br>anti Mouse | Goat | 1/1000 | Molecular Probes | A21125 |
|  | **Alexa Fluor 647**<br>anti Mouse | Donkey | 1/1000 | Molecular Probes | A31571 |
|  | **Alexa Fluor 647**<br>anti Goat | Donkey | 1/1000 | Molecular Probes | A21447 |

### Image analysis and quantification

For sprout number quantification, phase contrast images acquired after 12 (iPSC) or 21 (mESC) days of 3D- endothelial differentiation were employed. Sprouts were identified according to their morphology and quantified manually. Confocal immunofluorescence images from the same time points were used to manually categorize cells in the sprouts based on the following criteria: i) to show either functional or dysfunctional adherens junctions (VE-Cadherin channel); ii) to have junction-associated branched actin; iii) the presence of attenuated VE-Cadherin; iv) to display either straight or thick intercellular β-catenin.

For other image quantifications, semi-automatic macros designed with Image J/Fiji software [30] were applied to confocal Z-stack pictures. The number of stress fibers in WT and p73KO-iPSC cells after 12 days of differentiation was quantified by “Ridge Detection” plug-in [31], applying the parameters previous established [32], selecting the stress fibers in individual cells using the F-actin channel.

AMOT fluorescence intensity at cell periphery was quantified by drawing a Region of Interest (ROI) around cell periphery using VE-Cadherin channel (Total) and another ROI delimiting VE-Cadherin intracellularly (Cytoplasm). “XOR” option was used to subtract the cytoplasm selection from the Total region and to obtain the Periphery region. Amot Mean Fluorescence Intensity (MFI) was measured using AMOT channel. For YAP ratio quantification a Gaussian Blur filter (σ = 2) was applied to the DAPI channel and images were binarized using an automatic threshold. The “Analyse particles” plug-in was used to identify the number of cells in the image. Then, the cell region was drawn manually (ROI). “XOR” option was used to subtract the nuclei selection from the cell region and obtain the cytoplasmic region. The mean intensity of YAP at nuclei and cytoplasmic region was measured. Proliferation analysis was performed using “Analysed particles” option for DAPI and KI67 channels. For *in vivo* permeability quantification, blood vessels were selected using the IB4-channel which was binarized with an automatic threshold. Blood vessel selection was overlapped with TER119 staining and “clear” or “clear out” options were used for outside or inside particle quantification, respectively. “Analyse particles” option was used to count TER119-stained particles for each condition and particle percentage was calculated for each image.

### RNA Sequencing (RNA-seq) and Transcriptome Data Analysis

Whole transcriptome sequencing from two biological replicates per condition was carried out at the Genomics and Proteomics Service of the National Centre for Oncological Research (CNIO, Madrid). RNA-seq was conducted on the HiSeq Illumina and 20 million reads were generated for each library. Sequencing reads were aligned to the transcriptome with TopHat (v 2.1.1; http://ccb.jhu.edu/software/tophat/index.shtml) provided with known gene annotations and other transcript data (GTF file) taken from the Ensembl (release 84) gene set for the GRCm38/mm10 mouse genome assembly. Alignment data were quantitated with HTSeq [33]. Differential gene expression analysis was subsequently performed with DESeq2 [34]. Genes were assigned as differential expressed genes (DEGs) if they had an adjusted p-value < 0.05.

Gene ontology (GO) and Functional Annotation Clustering were performed using DAVID Bioinformatics Resources 6.834 [35, 36], choosing level 3 of GO terms specificity. Clustering results were plotted using the ggplot2 library from R (3.5.1) Core Team (2014). Genes shared by the selected clusters were identified, and the results were plotted with Cytoscape 3.7.2 [37]. Venn diagrams were created using BioVenn (http://www.biovenn.nl/) [38].

### RNA isolation, reverse transcription, and quantitative real-time PCR (qRT-PCR)

Total RNA isolation and cDNA synthesis were performed using the RNeasy Mini Kit (Qiagen, #74106) and the High-Capacity RNA-to-cDNA™ Kit (Applied Biosystems, #4387406), respectively. Gene expression levels were detected by qRT-PCR in a StepOnePlus™ Real-Time PCR System (Applied Biosystems) using FastStart Universal SYBR Green Master mix (Roche, #4913850001). All protocols were performed according to the manufacturer’s instructions. Primers sequences are listed in Table 2.

**Table 2.**
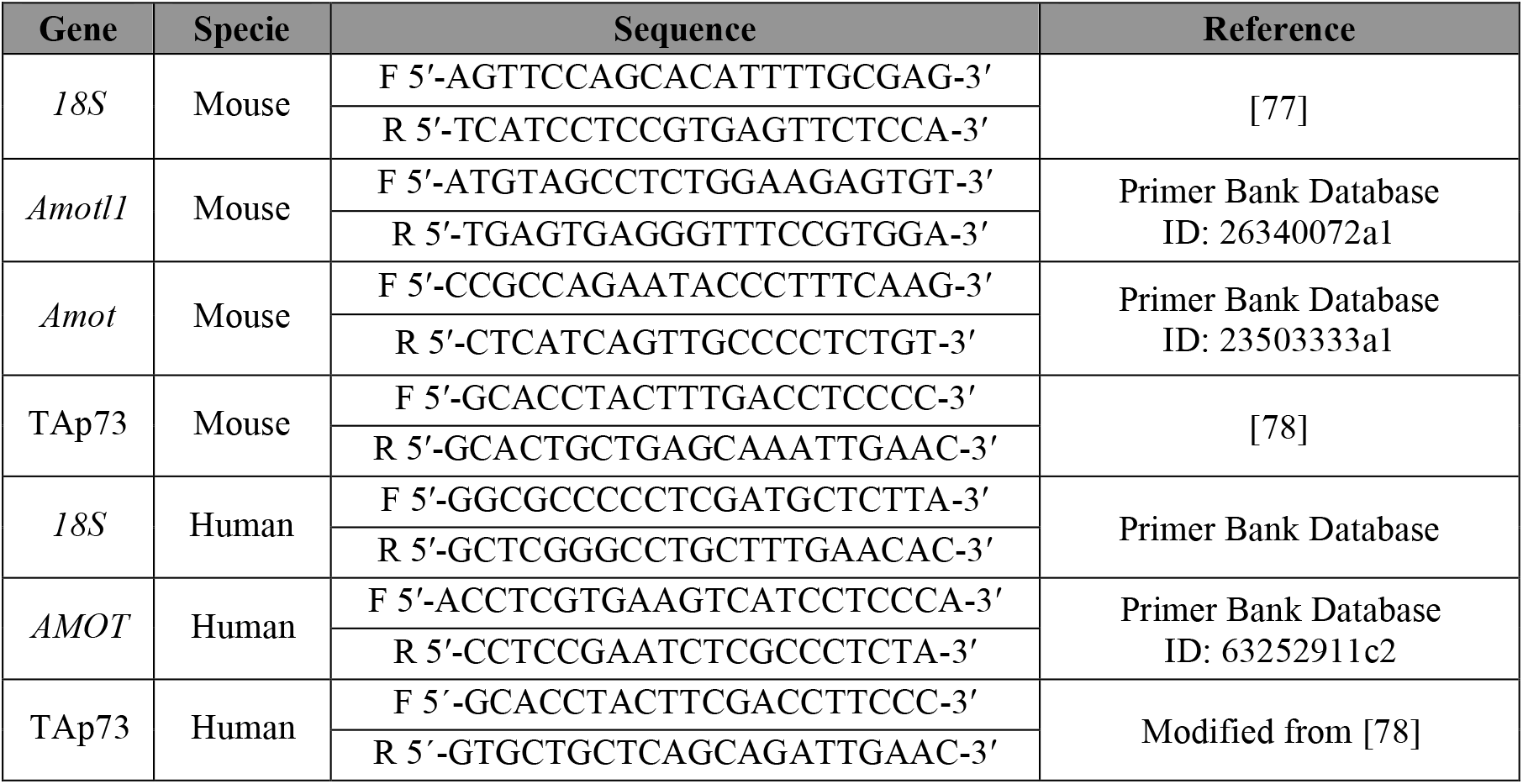
Primer sequences used for qRT-PCR.

### Chromatin immunoprecipitation assay (ChIP)

*In silico* prediction of the p73 responding element (RE) within the human *AMOT* gene (ENSG00000126016-; Transcript ID ENST 00000371959.7) was performed using the open access database for eukaryotic transcription factor binding profiles JASPAR [39]. Matrix models MA0861.1 and MA0525.1 were applied, and a site with a score of 13.3 was selected for further *in vitro* analysis. Primers encompassing the p73RE in human AMOT gene were as follows: F ⟶5’-TGTCCCCTTTTCTGCAGAGC -3′ and R⟶5’- GCTCCCCACTGACACGTTAA -3′.

ChIP analysis was carried out in HA-TAp73β-Saos2-Tet-On cells as described [25], using the following antibodies: anti-HA (Y-11) (Santa Cruz Biotechnology, #sc805), anti-p73 (EP436Y) (Abcam, # ab40658) and anti-p53 (C-19) (Santa Cruz Biotechnology, #sc1311).

### Western blot analysis

Protein isolation and electrophoresis were performed as previously published [22]. Nitrocellulose or PVDF (Polyvinylidene fluoride) membranes were incubated with rabbit anti- AMOT 1:600 (provided by Dr. Lars Holmgren, Karolinska Institutet, Stockholm, Sweden), anti-AMOT 1:500 (Cell Signalling, #43130S), anti-VE-Cadherin 1:1000 (BD, #610252), or mouse anti-GADPH 1:1500 (Abcam, #ab8245), diluted in 2.5% non-fat dried milk/TBS-T (Tris Buffer Saline-0.05% Tween 20), at 4 ºC o/n with gentle agitation. After washing with TBS-T, membranes were incubated with 25 ng/ml horseradish peroxidase-coupled secondary antibody (Pierce, #31430 or #31460) in 2.5% non-fat dried milk/TBS-T for 1 hour at RT. HRP-conjugated proteins were visualized with Super Signal West-Pico Chemiluminescent Substrate (ThermoFisher Scientific, #34087), followed by membrane autoradiography.

### Statistical analysis

Statistical analyses were performed using GraphPad Prism 9.3 software. Values were expressed as mean ± standard error of the mean (SEM). Differences were considered significant when *p* < 0.05 (\**p* < 0.05, \*\**p* < 0.01, \*\*\**p* < 0.001). Data distribution was analysed in all cases, and non-parametric tests were applied when necessary. In the latter scenario, statistical differences were calculated according to Mann-Whitney tests or to Kruskal-Wallis test with Dunn’s correction for multiple comparisons. For grouped data, a two-way analysis of variance (ANOVA) was carried out with Sidak’s or Tukey’s post hoc test to account for multiple comparisons. To test the association between the genotype and the categorical variables graphed in Figures 1-2, a contingency table analysis was used followed by Fisher’s exact test. In the case of YAP localization analyses, the ratio between YAP fluorescence intensity in the nucleus versus cytoplasm was calculated. For statistical reasons, a log transformation was applied to the data since the relative likelihood of sampling from a Gaussian versus a lognormal distribution was computed and the lognormal distribution was more likely.

**Figure 1.**
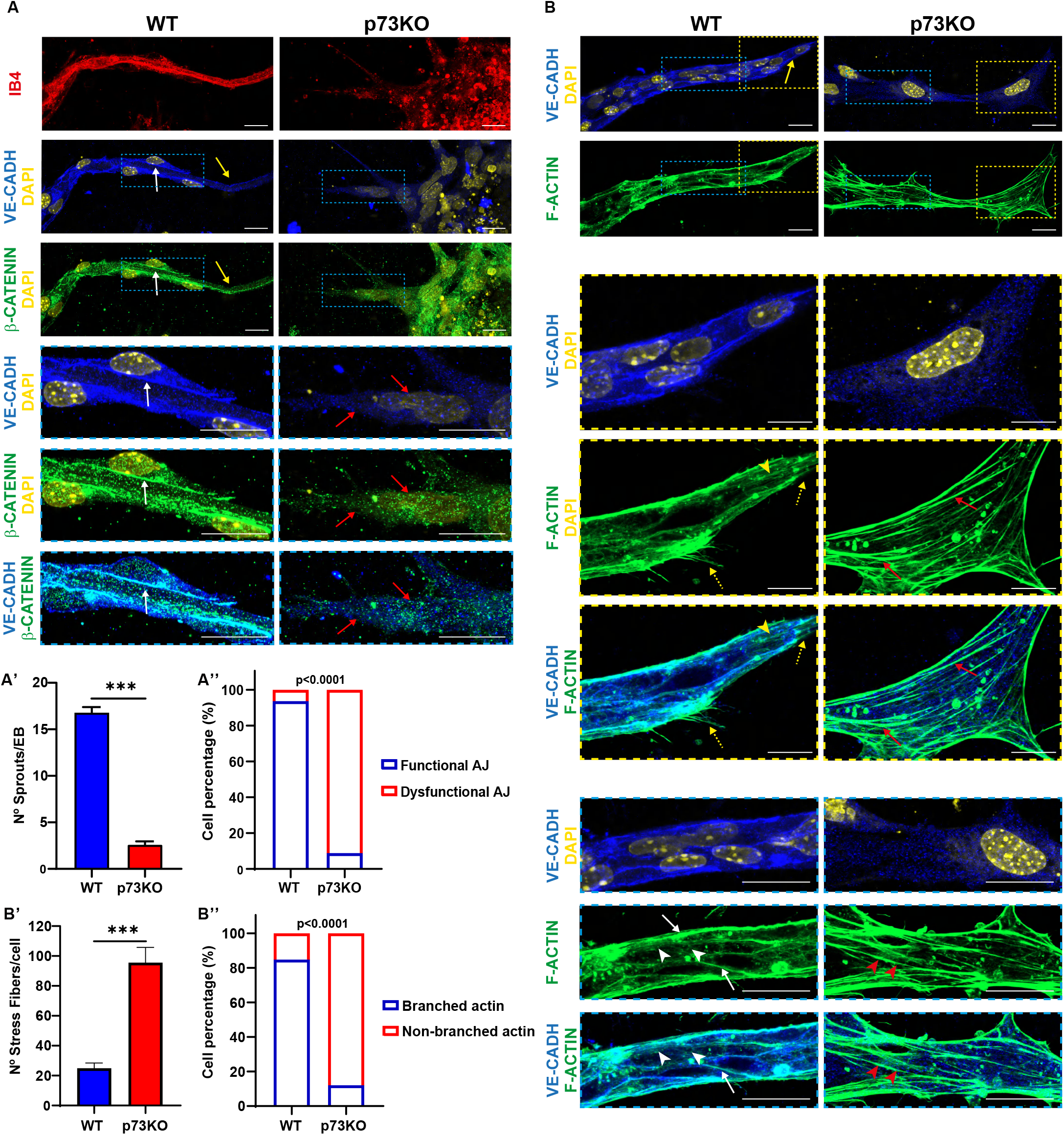
*Trp73* is required for the establishment of endothelial junctions during angiogenic sprouting *in vitro*. **(A, B)** Immunofluorescence (IF) analysis of endothelial adherens junctions (AJ) in IB4+ vascular sprouts from WT and p73KO-iPSC differentiated via EB formation and 3D culture conditions. After 12 days, sprouts were stained for VE-Cadherin (VE-CADH, blue) **(A, B)**, β-CATENIN (green) **(A)** and F-ACTIN (green) **(B)**. Nuclei were counterstained with DAPI (yellow). Scale bar: 20 µm. Magnifications of the areas marked with yellow or blue dashed squares are shown. Confocal microscopy images are representative of at least three independent experiments. **(A)** Yellow arrows indicate WT tip-cells, and white arrows point to stalk cells where VE-CADH staining is tightly co-localized with β-CATENIN. p73-deficiency results in altered junctional morphology independently of the cell position (red arrows). The number of sprouts per EB was quantified **(A’)** and cells in the sprouts were classified as showing either functional or dysfunctional AJ **(A’’)** depending on the VE-CADH staining (sharp and linear pattern *versus* scattered VE-CADH through the cell cytoplasm). At least 30 EB and 200 cells were quantified per genotype. **(B)** In WT sprouts, tip-cells (yellow dashed square) show thin filopodia (yellow dashed arrows) and stress fibers (yellow arrowheads), whereas in stalk cells (blue dashed square) VE-CADH interacts with F-ACTIN (white arrows) and AJs displayed junction-associated branched actin filaments (white arrowheads). Lack of p73 leads to a highly significant increase in stress fibers (red arrows and quantification in **B’**) and to the predominance of non-branched actin at cell junctions (red arrowheads and contingency analysis in **B’’**). Unpaired t-test was used to evaluate the differences in **A’** and **B’** (***p<0.001), whereas a Fisher’s exact test was used to compute the p value in the contingency analysis (p<0.0001).

**Figure 2.**
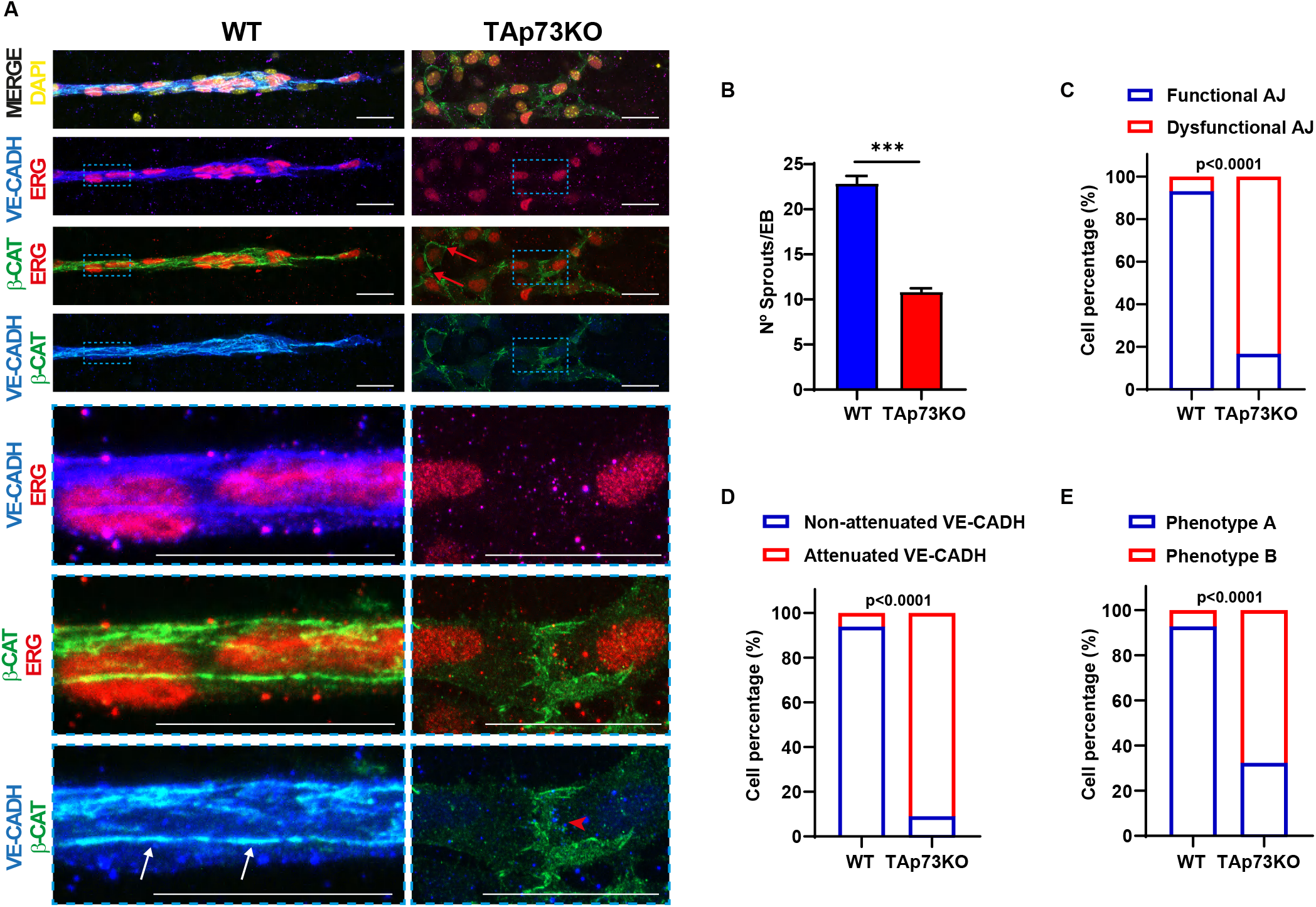
Endogenous TAp73 isoform is necessary for proper EC assembly and sprout formation during 3D-differentiation towards the vascular lineage. **(A-E)** Analysis of endothelial AJ in vascular sprouts from E14-WT and E14-TAp73KO mouse ESC. Cells were differentiated via EB formation and 3D-culture conditions and fixed after 21 days. **(A)** Immunofluorescence analysis of sprouts stained for VE-CADH (blue) and β-CATENIN (green). Nuclei were counterstained with ERG (red) and DAPI (yellow). Scale bar: 20 µm. Magnifications of the areas marked with blue dashed squares are shown. Confocal microscopy images are representative of at least three independent experiments. In WT-cells, EGR+ sprouts establish sharp AJ (white arrows). In the absence of TAp73, only some EGR+ cells establish AJ (red arrows), but the rest have thick intercellular accumulations of β-CATENIN and no proper VE-CADH staining (red arrowhead). **(B-E)** The difference in the number of sprouts between WT and TAp73KO EBs was quantified and cells within the sprouts were assigned by visual inspection into the following categories: **(C)** functional or dysfunctional adherens junctions according to the VE-CADH staining pattern; **(D)** attenuated or non-attenuated VE-CADH signal; **(E)** straight or thick intercellular β-catenin accumulation, named as Phenotype A or B respectively. At least 30 EB and 120 cells were quantified per genotype. Fisher’s exact test was used to assess if there was an association between the genotype and the qualitative variables defined above (p<0.0001).

## Results

### p73 is required for endothelial junction assembly during angiogenic sprouting and orchestrates transcriptional programs involved in cell-cell adhesion and blood vessel morphogenesis

To investigate the underlying mechanism by which p73 arranges the formation of vascular sprouts, we used a well-established angiogenesis model from iPSCs. Wild type (WT) and p73KO-iPSCs were differentiated to generate IB4 positive (IB4+) sprouts (Figure 1A). Endothelial cells in WT-sprouts established a front-rear polarity at the leading edge, where tip-cells adopted a finger-like shape with slightly low levels of VE-Cadherin that appears mainly cytoplasmic (Figure 1A and B, yellow arrows). These cells directed migration at the leading pole through the formation of thin filopodia (Figure 1B, yellow dashed arrows) and stress fiber formation (yellow arrowheads). Tip-cells polarized rear-ends were associated to stalk-cells via sharp and linear adherens junctions (AJ) (Figure 1A, white arrow). However, p73KO-iPSCs significantly developed less sprouts (Figure 1A’), and the few IB4+sprouts that were detected lacked front-rear polarity and had altered junctional morphologies (Figure 1A’’) independently of the cell position, with VE-Cadherin and β-catenin scattered through the cell cytoplasm (Figure 1A, red arrows). Moreover, cells showed an increase in the number of actin bundles forming stress fibers (Figure 1B, red arrows, and B’). Thus, p73-deficiency not only affected the polarization of the sprouts, but also their cytoskeleton. Indeed, WT stalk-cells (Figure 1B, blue dashed box) established AJ with a linear VE-Cadherin pattern that strongly interacted with F-actin (Figure 1B, white arrows) and displayed junction-associated branched actin filaments (white arrowheads), while the majority of cells in the p73KO-sprouts lacked branched actin at cell junctions (Figure 1B, red arrowheads, and B’’).

To address endogenous TAp73 requirement for proper EC assembly and sprout formation, we applied the same approach to mESCs with specific inactivation of the TAp73 isoform (E14-TAp73KO, [22]. Again, lack of TAp73 significantly impaired the formation of sprouts when compared to WT, but the effect was less dramatic than in p73KO-EBs and some ERG+ sprouts were found (Figure 2A and B). In WT-sprouts, EC established sharp AJ with linear VE-Cadherin/β-catenin colocalized staining (Figure 2A, white arrows). However, there was a highly significant association between TAp73 deficiency and the presence of sprouts with dysfunctional AJ (Figure 2C) and attenuated VE-Cadherin staining (Figure 2D) which appeared as dots scattered through the cytoplasm. Few p73KO-EC, mostly located at the base of the sprout, displayed a WT-like AJ with intercellular straight β-catenin staining (Figure 2A, red arrows, and Figure 2E, phenotype A), but the majority showed AJ with thick intercellular accumulation of β-catenin (Figure 2A, red arrowhead, and Figure 2E, phenotype B). These data demonstrate that TAp73 isoform is necessary for the establishment of EC junctions during sprouting angiogenesis.

To address whether TAp73 could modulate transcriptional profiles associated with the regulation of intercellular junctions, we compared the transcriptomes of undifferentiated p73KO-iPSCs transfected with an empty vector with those with TAp73 ectopic overexpression (+TAp73). The DESeq2 analysis identified 301 DEGs with a p- adj<0.05 (Supplementary Table 1). Functional annotation clustering of the 235 upregulated genes revealed the implication of these DEGs in biological processes (BPs) such as development, cell motility, blood vessel morphogenesis, cell-cell adhesion and actin organization, among others (Figure 3A and Supplementary Table 2). In particular, some of the GO terms with the higher fold enrichment were related to cell junction organization (GO:0034330, Fold enrichment= 5.97, FDR 1 × 10^−3;^), while the BPs with the highest significance corresponded to tissue development (GO:0009888, Fold enrichment=3.28, FDR 1.77 × 10^−17^) (Supplementary Table 2). Analysis from the clusters “cell-cell adhesion” and “blood vessel morphogenesis” to generate an enrichment map (Figure 3A’) showed that almost half of the DEGs (47%) in the “cell-cell adhesion” cluster were common to all the GO terms strongly associated to this cluster (FDR ranging from 1.65 × 10^−12^ to 3.09 × 10^−7^). Similarly, the BPs included in the “blood vessel morphogenesis” cluster shared one third of the DEGs (FDR ranging from 5.23 × 10^−9^ to 8.08 × 10^−3^), altogether suggesting that there is a preserved gene set which is central to the establishment of cell-junctions and the formation of blood vessels that is regulated by TAp73, all consistent with the reported TAp73 involvement in epithelial architecture [40].

**Figure 3.**
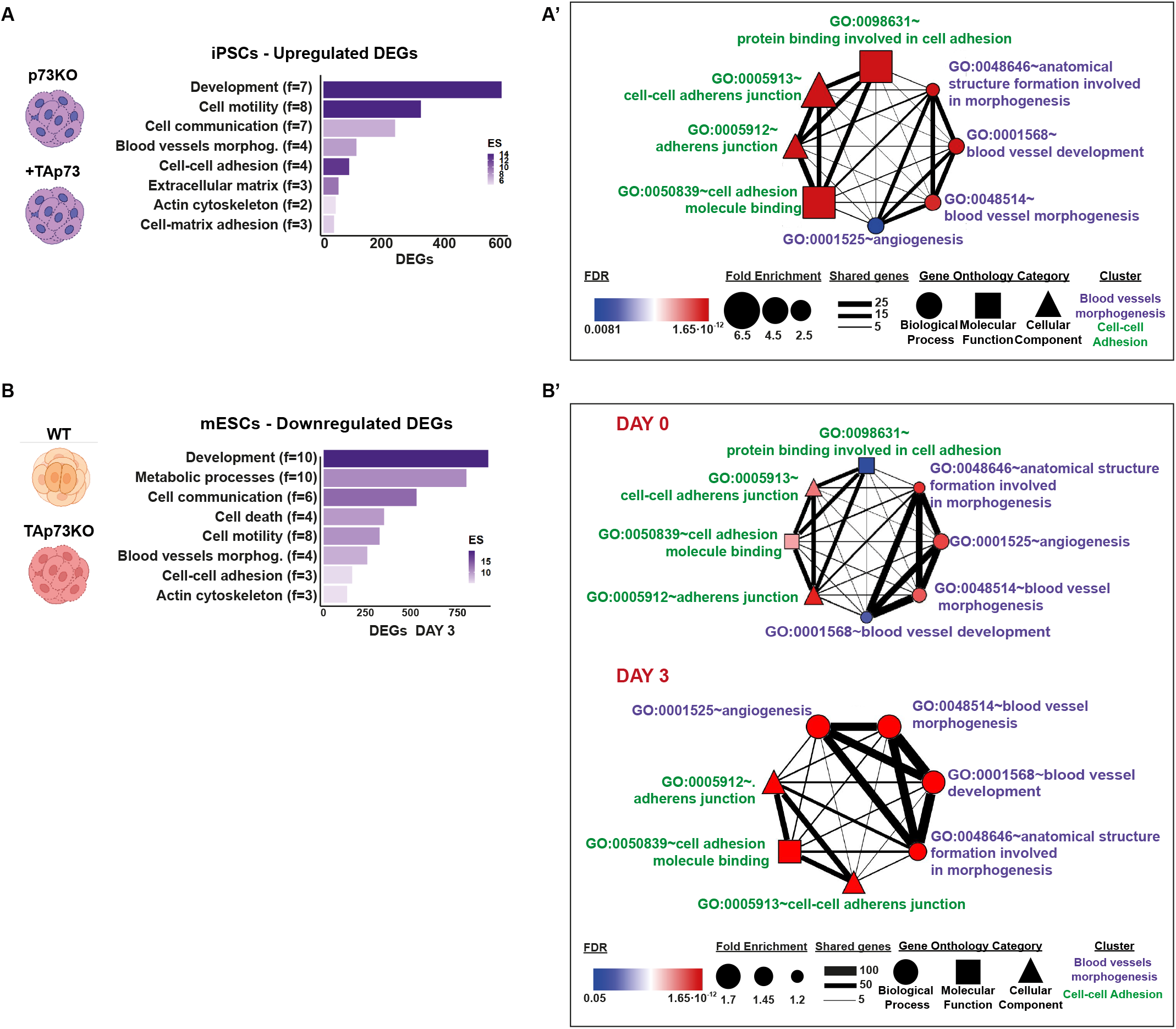
TAp73 modulates transcriptional programs involved in cell-cell adhesion and blood vessel morphogenesis. Transcriptomic studies were performed in p73KO-iPSCs after TAp73 ectopic overexpression (+TAp73) in comparison with cells transfected with an empty vector **(A, A’)**. RNA-seq data from previous studies (available at https://open.scayle.es/dataset/lopez-ferreras-et-al-cancers-2021) was also revisited **(B, B’)**. Differentially expressed genes (DEGs) with a p-adj<0.05 were used for GO analysis using DAVID. **(A, B)** Functional annotation clustering of up-regulated DEGs in p73KO-iPSCs after TAp73 re-expression **(A)** and downregulated DEGs in E14-TAp73KO cells under differentiation conditions-Day 3 **(B)**. The number of terms that compose each cluster is indicated (f) and the enrichment score is depicted by the color gradient. **(A’, B’)** Network analysis in iPSCs cells **(A’)** and mESCs (**B’**, Day 0 top panel, Day 3 bottom panel) of the following clusters previously identified: Cell-cell adhesion and Blood vessels morphogenesis (GO terms depicted in green and violet, respectively). Nodes in the graph represent the GO terms included in each annotation cluster. The node color indicates the false discovery rate (FDR) for the DEG association to each term, while the node size represents the Fold Enrichment. The edges connect overlapping terms, with the edge thickness depicting the magnitude of the overlap.

To further reinforce our observations and to help us to elucidate the molecular basis of p73 regulation of intercellular junctions and angiogenesis, we revisited a published RNA-seq study in mESCs, comparing parental E14TG2α cells to E14-TAp73KO, either under proliferation conditions (Day 0, D0) or after three days of EB-induced differentiation (D3), when endogenous TAp73 is known to be upregulated [22]. It has been reported that at D3 of EB formation, in the absence of LIF, the EBs contain a mixture of mesodermal and ectodermal progenitors and express mesendodermal marker genes including *Eomes, Foxa2* or *Gsc*-*Goosecoid* [41]. This stage is previous to the expression of other endothelial markers like *Pecam1* or *Tek* (*Tie-2*), reported to be rapidly upregulated after day 4 [42]. In our model, we detected at D3 a significant increase of some endothelial progenitor cell markers [43] including *Cd34* and *Etv2* [22]. In line with our previous results, clustering analysis of downregulated DEGs (Figure 3B) disclosed BPs significantly associated (FDR<0.05) with development, cell communication, cell-cell adhesion and blood vessels morphogenesis. Interestingly, the molecular interaction networks (Figure 3B’) confirmed that, concomitantly with the reported TAp73 upregulation from D0 to D3, the number of shared genes within the GO terms of the clusters, the fold enrichment and, particularly, the FDR increased at D3, altogether strengthening the idea of p73 as a regulator of the vascular architecture through the control of transcriptional programs involved in cell junction establishment.

### TAp73 regulates Angiomotin expression and downstream function

To identify possible p73-effector targets, we performed a comparative analysis using: i) the DEGs from the above- RNA-seq mentioned experiments in iPSCs and mESCs; ii) published transcriptomic data of upregulated DEGs after TAp73 expression in p73KO mouse epithelial granulosa cells [44]; and iii) a list of p73 putative targets identified by ChIP-seq [45]. Some of the overlapping genes between these studies (Figure 4A) were well-known p73-transcriptional targets, such as *Mdm2* [46, 47], *Cdkn1a* [48], *Itgb4* [49], or *Trim32* [50], backing up the validity of our models to identify TAp73 target genes. Interestingly, among the overlapping genes, we found the angiomotin family members *Amotl1* (with predicted p73 binding sites [44]) and *Amot*, not yet described as a p73 target. AMOT contributes to the assembly and stability of EC junctions [5], controls polarization of vascular tip-cells [7, 51] and integrates spatial cues from the extracellular matrix to establish functional vascular networks [52], altogether making AMOT an interesting candidate as a p73-downstream effector in the maintenance of EC junctional integrity.

**Figure 4.**
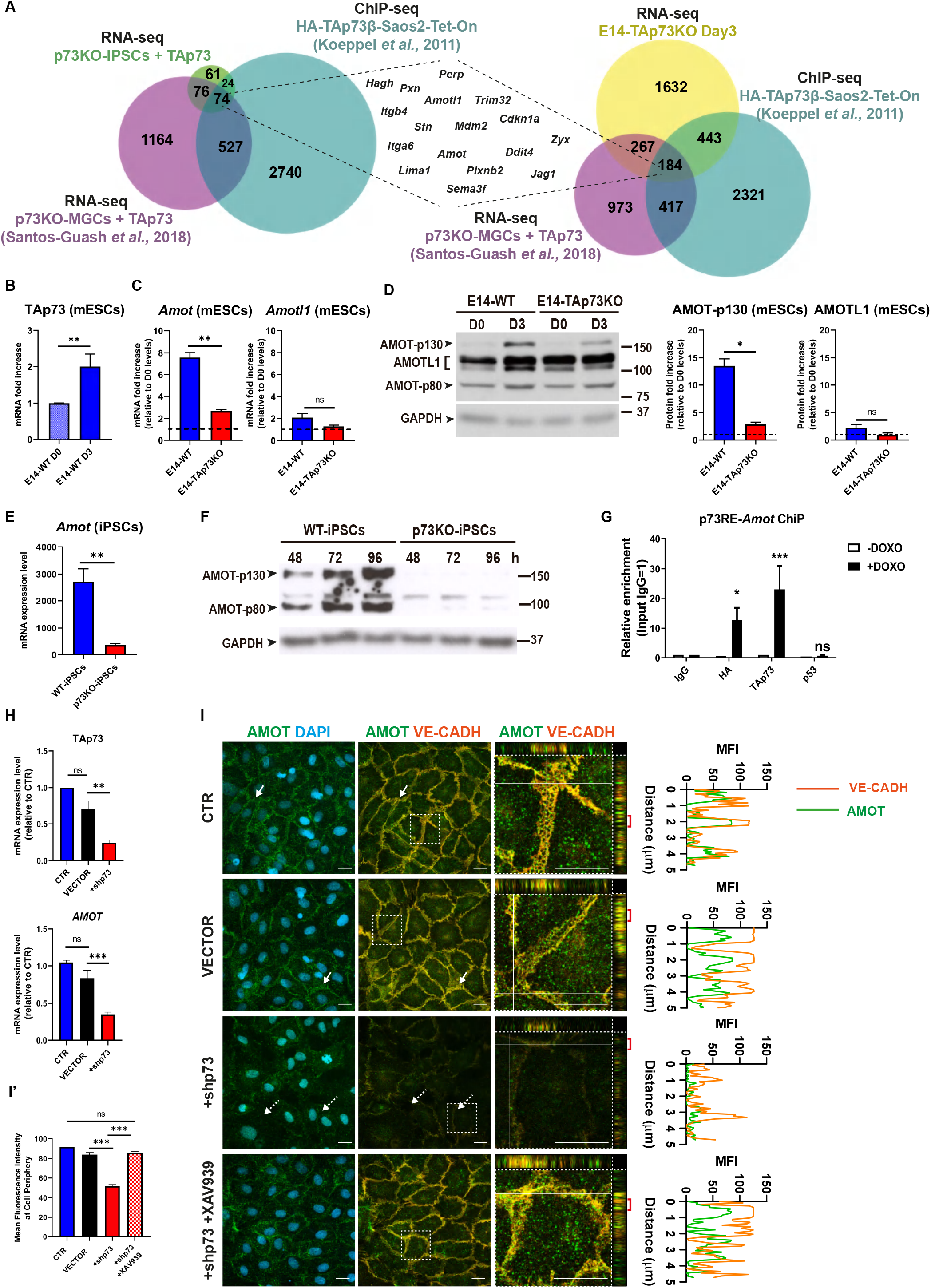
AMOT, a member of the Angiomotin family, is a direct TAp73 transcriptional target. **(A)** Comparison of the RNA-seq experiments in Figure 1 (upregulated DEGs in p73KO-iPSCs + TAp73 and downregulated DEGs in the case of E14-TAp73KO Day 3) with previously published transcriptomic data and ChIP-seq data from the indicated studies. **(B, C)** qRT-PCR analysis of TAp73 (B), or *Amot* and *Amotl1* expression in mESCs of the indicated genotypes under differentiation conditions. Values correspond to the fold increase relative to E14-WT Day 0 (B) or to their respective D0 values for each genotype (C). At least three independent experiments were performed. 18S Ribosomal RNA was used as housekeeping gene. **(D)** Western blot analysis and quantification of AMOT and AMOTL1 expression in mESCs of the indicated genotypes. Graphs shows the mean of two independent experiments yielding similar results. **(E, F)** qRT-PCR analysis (E) and western blot (F) were performed to evaluate *Amot* expression in WT- and p73KO-iPSCs. **(G)** ChIP analysis shows direct binding of TAp73 and HA-TAp73 to the selected p73RE in HA-TAp73β-Saos2-Tet-On cells following TAp73 induction by Doxycycline treatment (–/+DOXO). ChIP was performed using an isotype-control antibody (rabbit IgG) and anti- p53 as negative controls, or an anti-HA or anti-TAp73 antibodies for quantifying TAp73 binding. qPCR with primers specific to the p73-RE identified in the human *AMOT* promoter was performed and the fold enrichment relative to IgG precipitation is represented. **(H, I, I’)** p73 knockdown correlates with lower AMOT RNA and protein levels (H, I’, respectively) and a defective AMOT localization (I) in HUVEC. **(H)** qRT-PCR analysis of TAp73 and *AMOT* expression after p73KD with an shRNA targeting p73 (+shp73). Non-infected HUVEC (CTR) and cells transduced with a non-silencing empty vector (VECTOR) were used as controls. **(I)** Representative confocal micrographs of confluent monolayers from control cells and p73KD cells, with or without treatment with the tankyrase inhibitor XAV939. Cells were immunostained for AMOT (green) and VE-CADH (orange); nuclei were counterstained with DAPI (light blue). Magnifications of the areas marked with white dashed squares are displayed. Orthogonal projections of the magnified areas show AMOT localization at cell junctions colocalizing with VE-CADH. The histograms at the right represent the mean fluorescence intensity (MFI) profiles of AMOT and VE-CADH expression in the region indicated by the red bracket at the orthogonal projection. Scale bar: 20 µm. **(I’)** Quantification of AMOT mean fluorescence intensity at cell periphery is shown. At least 100 cells from 3 independent experiments were quantified per condition. In all panels, values represent mean ± SEM of three independent experiments. *p<0.05, **p<0.01, ***p<0.001, ns: non-significant.

Expression analysis of E14-WT mESCs during differentiation corroborated that, coinciding with an increase in TAp73 levels from D0 to D3 (Figure 4B), there was a strong upregulation of Amot expression at RNA and protein levels (Figure 4C and D, respectively, blue bars) and, to a much lesser extent, of Amotl1. This enhancement of Amot expression, but not of Amotl1, was significantly hindered in the absence of TAp73 (Figure 4C and D, red bars), suggesting that endogenous TAp73 plays a role in angiomotin upregulation during differentiation. Consistently, Amot levels were abated in p73KO-iPSCs (Figure 4E and F) and transfection of TAp73 in these cells induced AMOT expression 24 hours after the maximum TAp73 protein expression (Supplementary Figure 1A and 1B); then the effect subsided. Moreover, when using a well-known model for inducible TAp73 expression, like the HA-TAp73-Saos2-Tet-On cells, HA-TAp73-induction by doxycycline treatment prompted a significant increase in *AMOT* expression (Supplementary Figure 1C), suggesting that p73 regulation of this gene could occur in different cellular contexts.

To demonstrate that *AMOT* was a direct TAp73-transcriptional target, we searched *in silico* for p73 response elements (RE) within the *AMOT* proximal promoter and regulatory region and identified a strong p73RE (score: 13.3) located in intron 1. ChIP assays were carried-out in HA-TAp73-Saos2-Tet-On cells (Figure 4G). Crosslinked cellular extracts were immunoprecipitated (IP) using either an anti-TAp73 to detect all TAp73, or anti-HA antibodies for the induced HA-TAp73. As negative controls, we IP with an isotype immunoglobulin (IgG) or an anti-p53 antibody. Both anti-HA and anti-TAp73 pulldowns showed a significant enrichment compared to negative controls (Figure 4G), demonstrating that *AMOT* is a direct TAp73 transcriptional target.

In an attempt to move towards an EC model, we differentiated iPSC into IB4+EC and asked whether previously reported angiomotin expression during differentiation [53, 54] was p73-mediated (Supplementary Figure 1D, E). Comparison between AMOT levels in WT-EBs (P0) and IB4+EC in their first passage (P1) corroborated that, as reported, AMOT-p130 expression increased with differentiation, and even though the levels decreased in successive passages (P1 vs P3), they were always significantly lower in p73-deficient EC, independently of passage number (Supplementary Figure 1D).

We further investigated the need of p73 function for AMOT proper expression using the HUVEC endothelial cell model (Figure 4H, 4I). TAp73 levels are low in HUVEC; nevertheless, silencing of *TP73* gene with short hairpin RNA (+shp73, p73KD) was robust compared to the non-infected control (CTR) and the empty vector-transduced cells (VECTOR). p73KD significantly affected AMOT expression at RNA and protein levels (Figure 4H, I, I’) resulting in a diminished expression at the plasma membrane. In addition, orthogonal projections of confocal images demonstrated that in CTR and VECTOR cells, AMOT was detected at the intercellular junctions, together with VE-Cadherin (Figure 4I, white arrows and orthogonal projections), but this localization was largely lost in +shp73 cells (dashed white arrow, and orthogonal projection side quantification). AMOT levels and the expression pattern could be reinstated by treatment of +shp73 HUVEC with XAV939, a tankyrase inhibitor that is known to efficiently stabilize AMOT [55] (Figure 4I, I’ and Supplementary Figure 2). Thus, the change of expression pattern (less AMOT at intercellular junctions) could be a consequence of AMOT lower expression and/or its redistribution to another cellular compartment.

It caught our eye that VE-Cadherin levels in p73KD cells appeared to decrease at the intercellular junctions, while AMOT stabilization seems to recuperate them (Figure 4I). It has been reported that the C-terminal tail of VE-Cadherin is removed during endocytosis or subsequent trafficking of the cadherin through the endosomal system and that the majority of this internalized cadherin failed to be labelled with a C-terminus antibody [56], like the one used in our study. Thus, to address whether the disappearance of VE-Cadherin from intercellular junctions after p73KD was caused by alterations in protein traffic or by protein loss, we analyzed total protein levels by western blot. Our data detected a very small decrease of total VE-Cadherin upon p73KD, which was significantly recovered upon AMOT stabilization (Supplementary Figure 2), suggesting that the observed effect was most likely due to the alteration of VE-Cadherin subcellular distribution, rather than protein loss.

Next, we addressed whether p73 regulation of *AMOT* impinged on its downstream functions. Even though it is not yet clear how AMOT acts to promote angiogenesis *in vivo* [52], one of its reported scaffolding functions is the cytoplasmic sequestration of YAP [57]. YAP promotes barrier function maintenance [58] and gets relocated to the nucleus upon disruption of cell junctions [59]. Thus, we used YAP nuclear/cytoplasmic ratio as a readout of AMOT function. Confocal microscopy analysis in non-treated control cultures showed EC with frequent YAP nuclear exclusion (Figure 5A, yellow arrowheads) and similar nuclear/cytoplasmic ratios between CTR and VECTOR cells (right graph). Downregulation of p73 biased YAP distribution toward the nucleus (Figure 5A, yellow arrows) as reflected by a significantly higher YAP nuclear/cytoplasmic ratio, indicating that lack of p73 altered YAP localization in EC. Treatment of +shp73 cells with XAV939 recovered AMOT junctional localization (Figure 5A, white arrows) and correlated with an increase in YAP nuclear exclusion (dashed yellow arrows) and a significant decrease of the nuclear/cytoplasmic ratios to those similar to controls (right graph), altogether demonstrating that the attenuated AMOT levels were accountable for the observed changes. Moreover, under these conditions, orthogonal projection analysis of boxed areas in Figure 5A revealed that stabilized AMOT colocalizes with YAP at the plasma membrane as well as in the cytoplasm (Supplementary Figure 3, white and blue arrows, respectively). AMOT acts as a scaffold protein to promote localization of YAP to the cytoplasm, cell junctions and actin cytoskeleton [60-63]. In particular, AMOT mediates contact inhibition by directly binding and sequestering YAP at the plasma membrane and cytoplasm [57]. Thus, we sought to address if p73-downregulation would affect YAP sequestration triggered by cell-cell contact. In sparse cultures, YAP protein was predominantly localized in the nucleus in controls and +shp73 HUVEC (Figure 5B and C, yellow arrows), with cells showing YAP nuclear/cytoplasmic ratios over 1.68 (Figure 5B, right graph) which correlated with a high percentage of proliferating cells (Figure 5C, right graph). As expected, in confluent control cultures many cells displayed YAP nuclear exclusion (Figure 5B and C, white arrows) and lower YAP nuclear/cytoplasmic ratio, all associated with a strong decrease in proliferating cells (Figure 5C). Conversely, in +shp73 EC under the same confluency conditions, we detected little nuclear exclusion (yellow arrows) and the nuclear/cytoplasmic ratio barely changed between the two seeding conditions (low [1.57 mean ratio] vs. high-density [1.46 mean ratio]). Moreover, AMOT/YAP-dependent contact-inhibition of cell proliferation was abated in the absence of p73 (Figure 5C).

**Figure 5.**
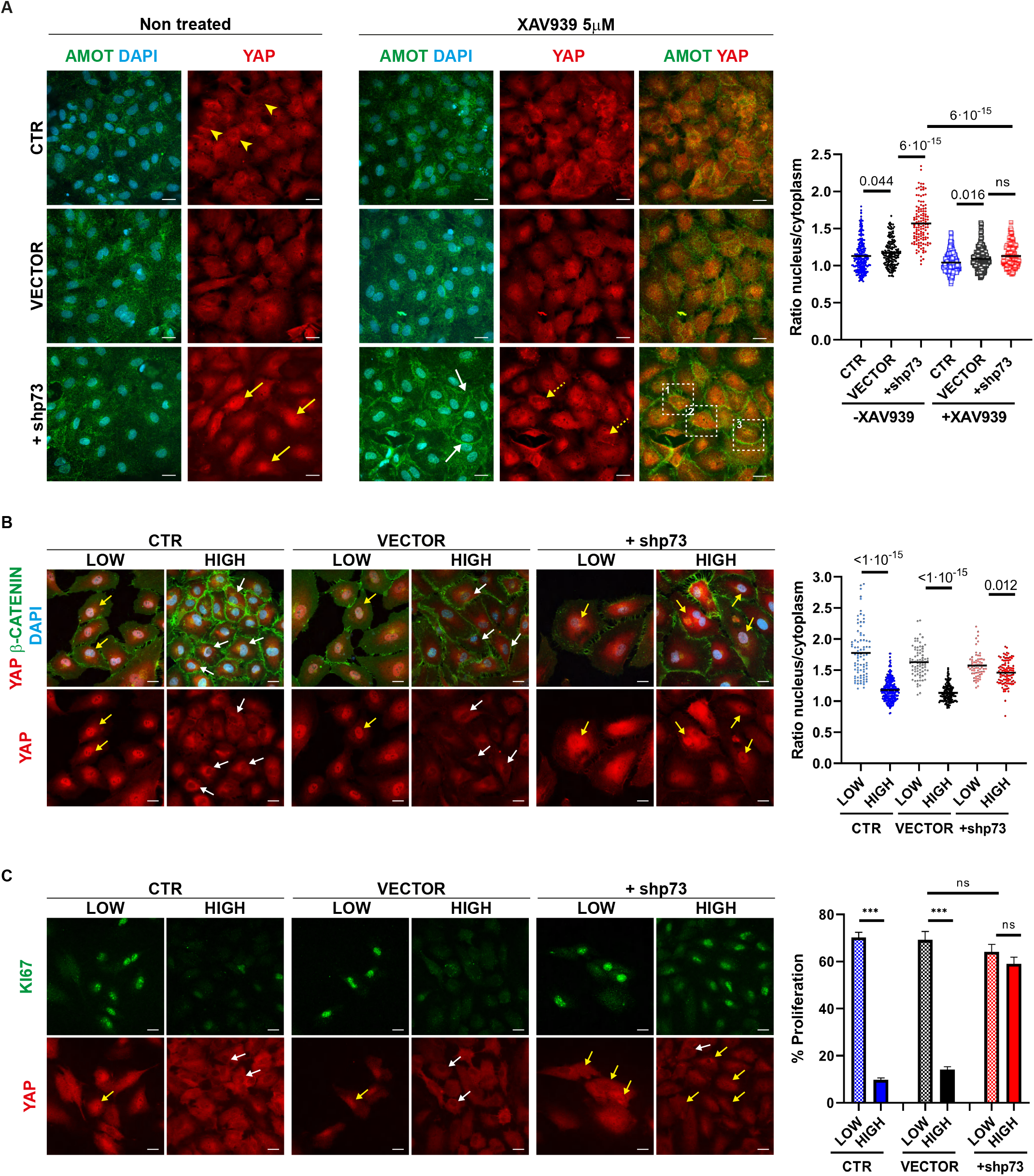
p73 regulation of AMOT impinges on its downstream functions and controls AMOT-mediated YAP sequestration during cell contact inhibition. **(A)** Non-infected control HUVEC (CTR), and cells transduced with a non-silencing empty vector (VECTOR) or with an shRNA targeting p73 (+shp73) were seeded at 7,5 × 10^4^ cells/cm^2^ and after 48-72 h, cells were treated with XAV939 for 24 h. Non-treated control cultures display frequent nuclear exclusion (yellow arrowheads) of YAP staining (red), whereas +shp73 cultures show a biased YAP distribution toward the nucleus (yellow arrows). Immunofluorescence analysis demonstrate that XAV939 treatment restores AMOT (green) levels and localization at intercellular junctions (white arrows) and correlated with a change in YAP localization in p73KD cells upon treatment (dashed yellow arrows). Orthogonal projections of the areas indicated by white dashed squares (1, 2, 3) are provided in Supplementary Figure 2, showing that stabilized AMOT colocalizes with YAP at the plasma membrane as well as in the cytoplasm. (**A**, right graph) Quantification of YAP nuclei/cytoplasm ratio after p73KD +/- XAV939 treatment. For these analysis, three independent infections were performed and at least five pictures from each experiment were randomly selected and quantified (n>100 analyzed cells across the three independent experiments). The obtained data followed a logarithmic distribution; thus, for the statistical analysis, a log transformation was applied to the data. The computed P value is indicated in the graph. **(B, C)** Analysis of YAP (red) localization and proliferation (**C**, KI67, green) in HUVEC at different seeding density conditions (Low - 1 × 10^4^ cells/cm^2^ versus High density - 7,5 × 10^4^ cells/cm^2^/High). Cell delimitation was discerned by β-CATENIN (**B**, green) staining. Nuclei were counterstained with DAPI (**B**, light blue). Scale bar: 20 µm. Under sparse culture conditions (low density) YAP localized mainly at the nucleus (yellow arrows), either in controls or p73KD cells. Upon contact inhibition (high density) many CTR and VECTOR cells displayed YAP nuclear exclusion (white arrows); however, p73KD cells (+shp73) showed little or no nuclear exclusion (yellow arrows) upon confluency. Quantification of YAP nuclei/cytoplasm ratio was performed as in (A) (n>60 cells for Low-density and n>100 for High-density condition). **(C)** Quantification of the proliferation percentage after p73KD (right panel). Bars represent mean ± SEM of three independent experiments. ***p<0.001, ns: no significant.

These observations place p73 as a novel regulator of the AMOT/YAP contact-inhibition pathway, essential for the fine tuning between junctional stability and angiogenic activation of ECs.

### p73 regulates endothelial junction dynamics via AMOT

Next, we asked whether TAp73 controls the establishment of EC junctions, at least in part, through the regulation of AMOT. Confirming our previous observations, confocal microscopy analysis of +shp73 HUVEC confluent cultures revealed a strong alteration of AJ, visualized by VE-Cadherin staining (Figure 4I and 6A). To accurately describe those differences, we took advantage of previous studies that correlated AJ morphology with cellular activities [64]. We observed that CTR and VECTOR cells frequently showed thick/reticular and reticular VE-Cadherin staining at the cell junctions (Figure 6A, pink and green arrow, respectively). They also establish straight and thick junctions (purple and orange arrows) that resemble the stalk-cell behavior [64]. However, interrupted serrated junctions (yellow arrows, VE-Cadherin fingers herein) were present in lesser proportion in controls. Strikingly, p73KD modified the distribution of junctional categories resulting in a sharp increase of fingers (Figure 6A, yellow arrows and bars). This type of interrupted junctions has been associated with actively rearranging cells and augmented permeability in confluent monolayers [58, 65, 66]. Furthermore, p73KD caused a highly significant decrease in linear (straight and thick) and reticular junctions (purple, orange, and pink and green bars, respectively), which are required to maintain junction integrity [67]. In line with this, p73KD led to junctional breaks in the monolayer (Figure 6A, white arrows). Analysis of Tight junctions (TJ) by ZO-1 expression also revealed an interrupted ZO-1 staining pattern, reflecting the accompanying alteration on TJ upon p73KD (Supplementary Figure 4).

**Figure 6.**
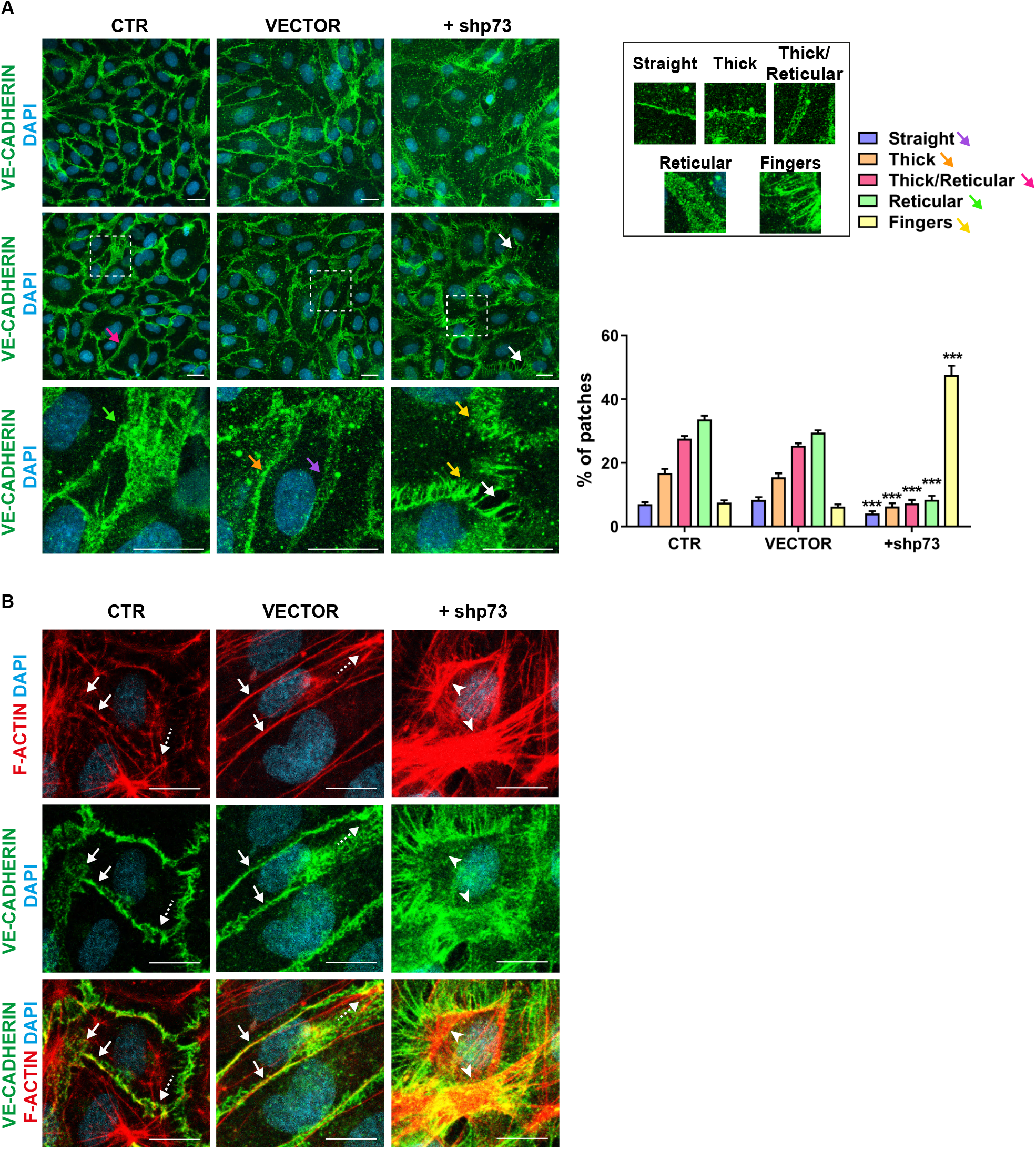
p73 regulates adherens junction and actin organization *in vitro*. **(A, B)** Representative confocal microscopy images of confluent HUVEC monolayers showing the alterations of adherens junctional morphologies after p73KD. Non-infected control HUVEC (CTR), and cells transduced with a non-silencing empty vector (VECTOR) or with an shRNA targeting p73 (+shp73) were stained for VE-CADH **(A, B**, green) or F-ACTIN **(B**, red). Nuclei were counterstained with DAPI (light blue). Scale bar: 20 µm. **(A)** Morphological analysis of VE- CADH labelled cell junctions in HUVECs after p73KD. Cell junction morphology analysis was performed based on previous publications [58]. Briefly, five morphological categories were defined: straight junctions (purple arrow), thick junctions (orange arrow), thick to reticular junctions (pink arrow), reticular junctions (green arrow) and fingers (yellow arrows). White arrows point to cellular gaps. Each image was divided in 36 patches and a morphological category was manually assigned to each patch. Representative patches (right panel) used for morphological classification are shown. Bottom images correspond to the magnifications of areas marked with white dashed squares. Bars represent the mean percentage ± SEM for each type of morphology Three independent transductions experiments were performed and ten pictures per experiment were randomly selected and classified (*n*=30 pictures per condition). p values indicate the statistical differences between +shp73 and VECTOR values (***p<0.001). There were no significant differences between CTR and VECTOR. **(B)** In control cells, cortical actin bundles (labelled in red) associate to VE-CADH (green) in linear and reticular junctions (white arrows), whereas the discontinuous junctions form stellate arrangements at cell borders (dashed white arrows). p73KD results in an increase of actin bundles appearing as stress fibers (white arrowheads).

In resting EC controls, cortical actin bundles associated with linear and reticular VE-cadherin junctions (Figure 6B, white arrows) are known to be critical for the regulation of barrier function [67]. In these monolayers, the discontinuous serrated junctions can be associated with stress fibers [68] that form actin stellate arrangements at cell-cell borders (dashed white arrow). These junctional fibers are increased in response to permeability-enhancing factors, such as TNF [68]. In accordance with the enhanced finger-like junctions in p73KD cells, we observed an increase in actin bundles appearing as stress fibers in these cells (Figure 6B, white arrowheads). These observations demonstrate that p73 is involved in the remodeling of endothelial junctions and the associated cytoskeleton and is required for the formation of reticular junctions and the repression of VE-Cadherin fingers formation, altogether potentially leading to defects in paracellular permeability.

To demonstrate AMOT requirement for p73 regulation of EC junction dynamics, we attempted to rescue p73KD- junctional defects by stabilizing endogenous AMOT protein. As before, XAV939 treatment of p73KD cells increased AMOT levels, returning AMOT to its colocalization with VE-Cadherin and ZO-1 and reshaping the morphology of AJ and TJ (Figure 7A, yellow arrows, and Supplementary 4, respectively). Furthermore, morphometric analysis demonstrated that AMOT stabilization significantly recuperated the formation of reticular junctions and reduced the formation of fingers in the treated p73KD HUVEC monolayers (Figure 7A, lower panel). This restoration of linear and reticular junctions is concomitant with actin reorganization that now appear as cortical actin bundles like in the controls (Figure 7B). Altogether these data unequivocally demonstrate that the stabilization of AMOT in p73KD HUVEC restored endothelial junctional dynamics and integrity; therefore, p73 regulates intercellular junction dynamics, at least in part, through the regulation of AMOT.

**Figure 7.**
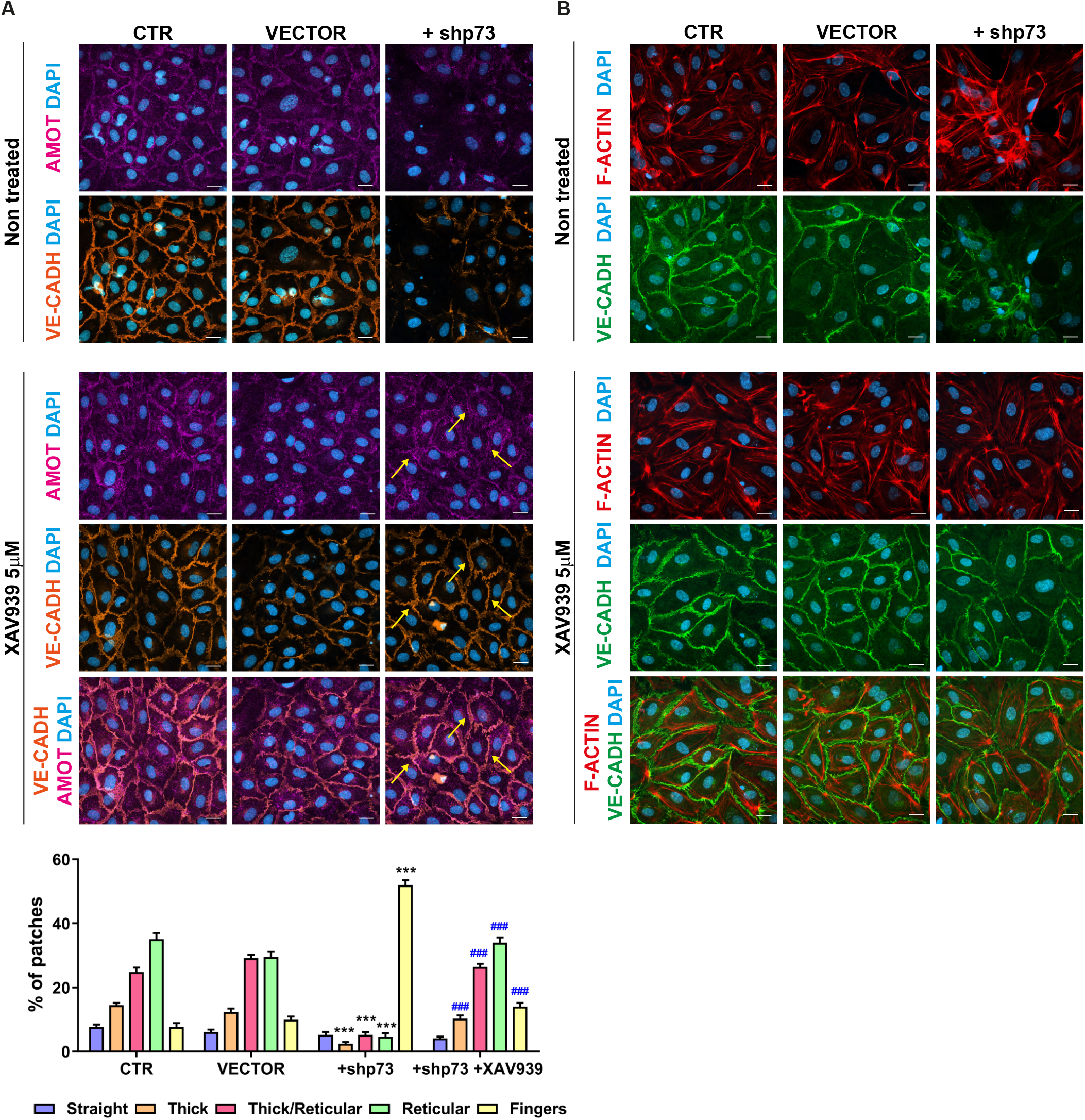
AMOT is a p73-downstream effector for the regulation of endothelial junctions and actin dynamics. **(A, B)** Confocal microscopy analysis of HUVEC after p73 knockdown and XAV939 treatment. Stabilization of endogenous AMOT levels (**A**, pink) by XAV939 treatment rescues the morphology of endothelial AJ as shown by VE-CADH staining (**A**, orange, yellow arrows; **B**, green) and the organization of F-ACTIN (**B**, red). Nuclei were counterstained with DAPI (blue). Scale bar: 20 µm. **(A**, low panel) Quantitative analysis of VE- cadherin cell junction morphology. Bars represent mean ± SEM (n=30 pictures per condition across three independent experiments). p values indicate the statistical differences between +shp73 and VECTOR values (***p<0.001) and between +shp73 untreated and XAV939-treated cells (###p<0.001). There were no significant differences among the morphological types between CTR and VECTOR.

### p73 is required for vascular barrier integrity *in vitro* and *in vivo*

To demonstrate the physiological relevance of p73 regulation of EC junctional dynamics we addressed whether p73-downregulation in EC translated into functional defects. Analysis of transendothelial permeability to dextran molecules revealed a significant increase in permeability in p73KD cells (Figure 8A). In addition, HUVEC monolayer barrier integrity was monitored in real-time by transendothelial electrical resistance (TEER) measurements (Figure 8B) and a significant decrease in TEER was observed in p73KD cultures. ECIS mathematical modelling of the TEER analysis yielded the basolateral membrane resistance values (Rb), Alpha values (center) and Cell membrane capacitance. TEER results showed that p73KD significantly reduced cell membrane resistance (Rb) values, which inversely correlates to paracellular permeability (Figure 8B). However, it did not affect alpha values, which reflect cell-matrix interactions, neither cell membrane capacitance, suggesting that the main effect of p73 downregulation, under the tested cell culture conditions, is the disruption of cell-cell junctions.

**Figure 8.**
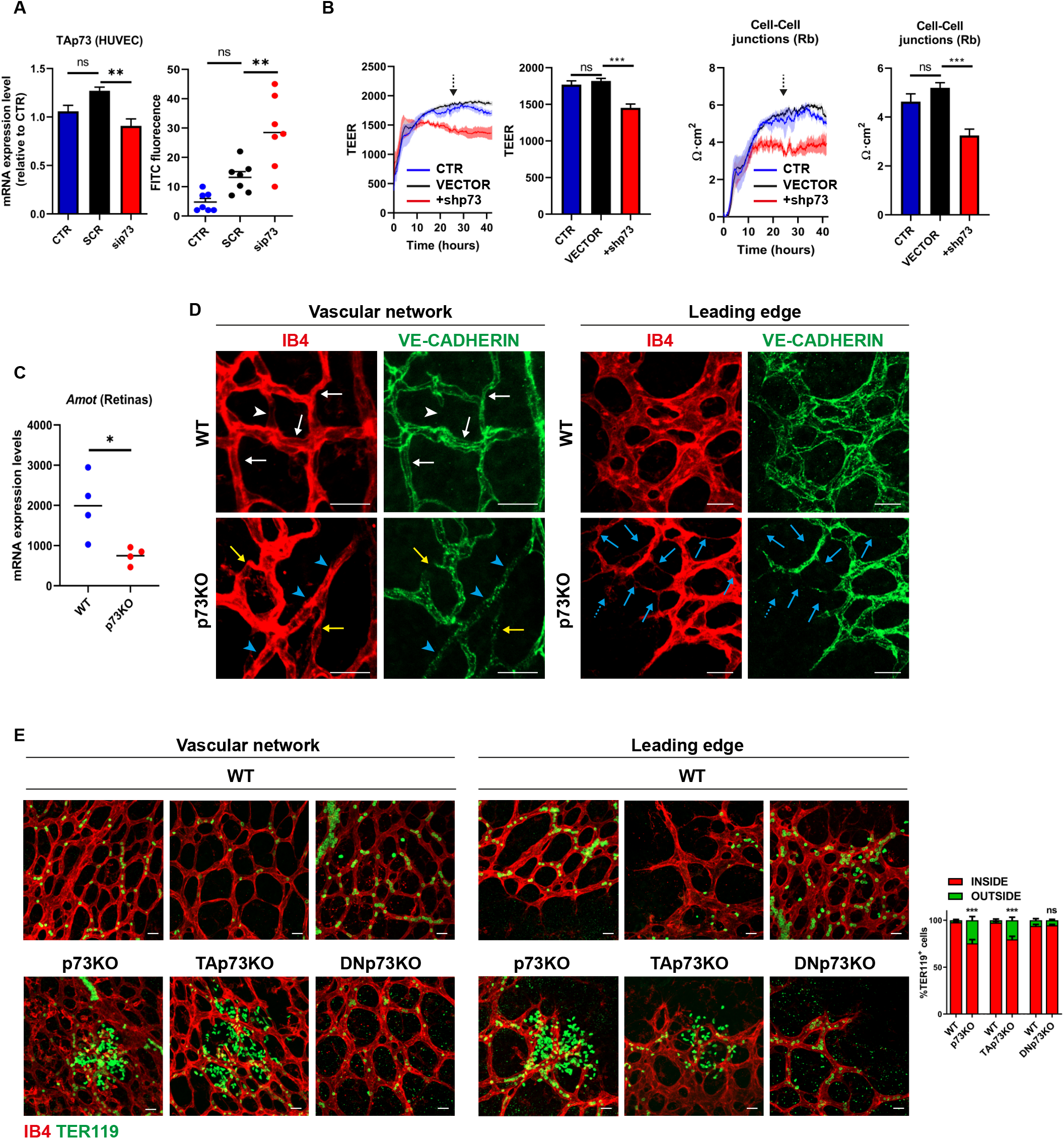
p73 is necessary for endothelial barrier integrity *in vitro* and *in vivo*. (**A**) Permeability assays to 250 kDa fluorescent dextran molecules in HUVEC monolayers comparing non-interfered (CTR), scrambled control (SCR) and p73-knockdown cells by siRNA (sip73). Cells were nucleofected and subsequently plated on the insert well. After 96 h, each insert was transferred to a fresh well plate and FITC-Dextran was added and incubated for 20 min at RT. Fluorescence (485 nm excitation/535 nm emission) was quantified in a plate reader. TAp73 expression level was analyzed by qRT-PCR in each case (left graph). (**B**) ECIS analysis of the effect of p73KD (+shp73) by TEER (Trans-endothelial electric resistance). The electric parameters were measured for 48 h after plating the cells, which had been transduced with the indicated lentivirus 3 days ahead. The 48 h-time course of TEER and of the basolateral membrane resistance values (Rb) are plotted, and graphs with the statistical analysis at the 24 h-time point (pointed by an arrow) are displayed. Three independent experiments were performed. (**C**) *Amot* mRNA expression levels in WT and p73KO mice retina analyzed by qRT-PCR. **(D, E)** Immunofluorescence analysis of P7 WT, p73KO, TAp73KO **(E)** and DNp73KO **(E)** mice whole mount retinas stained with IB4 (red), anti-VE-CADH (green) **(D)** and anti-TER119 (green) **(E). (D)** Representative confocal microscopy images of vascular networks and vascular leading edge in WT retina show linear intercellular junctions (white arrows) with unicellular segments lacking VE-CADH (white arrowhead). On the contrary, lack of p73 results in retina vessels with interrupted staining (yellow arrows), even in multicellular vessels (blue arrowheads), and in aggregations of IB4+EC (blue dashed arrows) close to sprouts with very little VE-CADH (blue arrows). Retinas from at least 3 mice per genotype were analyzed. Scale bar: 20μm. **(E)** Vessel integrity was assessed by quantifying the percentage of erythrocytes (TER119+cells) inside or outside the IB4+blood vessels (right panel). At least three mice per genotype were considered, and a minimum of 20 pictures were quantified for each genotype. Bars represent the mean percentage ± SEM. p value was calculated with respect to the WT littermates for each genotype. *p<0.05 **p<0.01, ***p<0.001, ns: no significant.

Finally, to ask if p73 and its isoforms play a physiological role in the maintenance of barrier integrity *in vivo*, we used mouse retinas to model physiological angiogenesis [69]. In retinal vessels, *Amot* is ubiquitously expressed at post-natal day 7 (P7) and is known to be essential for endothelial tip cell migration [52]. We detected significantly lower levels of *Amot* in the absence of p73 in P7 whole retina extracts by qRT-PCR (Figure 8C). VE-Cadherin staining in whole-mount retinas from WT and p73KO mice revealed profound differences in the arrangement of EC. Whereas in WT-retinas, VE-Cadherin delineates junctions that were thin, and mostly linear and continuous in all IB4+ vessels (Figure 8D, white arrows), in p73KO-retinas the EC junctions were tortuous with interrupted VE-Cadherin staining (yellow arrows). In WT-vessels we detected few unicellular segments lacking VE-Cadherin (Figure 8D, white arrowhead), while in p73KO-retinas there were many IB4+segments lacking VE-Cadherin staining (yellow arrows), even in multicellular vessels of normal caliber (blue arrowheads), indicative of defects in the establishment of multicellular tubes [17]. On the leading front of the retinas, the defective p73KO sprout morphology frequently correlated with aggregations of IB4+EC close to sprouts with very little VE-Cadherin (Figure 8D, blue arrows), arguing that the migration and/or the rearrangement of EC are perturbed in p73-defective vessels, and suggesting that vessels may have failed to anastomose or stabilize connections following sprouting. These defects in vessel morphology were coupled to defects in function, since p73KO and TAp73KO retinas, but not DNp73KO, exhibited large hemorrhages at the angiogenic front and at the vascular network in the remodeling retina plexus (Figure 8E). Altogether, our data strongly demonstrate that p73 plays a predominant role in maintaining the endothelial barrier function and that TAp73 is necessary for the maintenance of junctional integrity *in vivo*, regulating the rearrangements of EC whilst preventing bleeding in developing vessels.

## Discussion

The present work contributes to the landscape of p73 function in angiogenesis by presenting a comprehensive study in healthy endothelial cells which express physiological *TP73* levels. Our results provide an understanding of the mechanism by which p73 regulates intercellular junction dynamics in ECs, being essential for the maintenance of the endothelial barrier.

Endothelial junctions play a coordinating role in sprouting angiogenesis by polarizing the activity of the migration machinery and maintaining sprout cohesion [2]. Using 3D-endothelial differentiation models, we show that lack of p73, and TAp73 in particular, results in a strong impairment of the formation of endothelial sprouts and the establishment of AJ, demonstrating TAp73 requirement for these processes. Our *in vivo* data also support this scenario, since DNp73 was dispensable for the integrity of endothelial junctions in mice retinal vasculature, while lack of TAp73 led to functional defects. Nevertheless, specific elimination of TAp73 caused less dramatic defects than the total elimination of *TP73*, confirming the requirement of DNp73 pro-angiogenic function for a complete vascular morphogenesis process [9].

Even though TAp73 and DNp73 could have proangiogenic functions, it has been proposed that there may be differences in the mechanism by which they exert their activation and promotion of angiogenesis [70]. In agreement with the proposed function of p73 as a tissular architect [40] and supporting our hypothesis that p73 exerts a role on sprouting angiogenesis through the regulation of cell junctions, GO analysis of p73KO-iPSCs +TAp73 revealed that TAp73 orchestrates transcriptional programs involved in the regulation of cell-cell adhesion, developmental morphogenesis, and blood vessel morphogenesis. Similar results were obtained when revisiting an RNA-seq study in mESCs, comparing parental E14TG2α cells to E14-TAp73KO cells [22]. A comparative analysis between several published transcriptomic data and a list of p73 putative targets identified by ChIP-seq led us to focus our interest on AMOT, a member of the angiomotin family which is required for proper TJ and AJ activity [7], therefore contributing to preserve the sprout cohesion [4, 6, 7] and vascular permeability [5]. We identified *AMOT* as a direct TAp73 transcriptional target and showed that p73KD not only impinged in *Amot* expression, but also hindered its downstream functions, like the capacity to sequester YAP at intercellular junctions at the plasma membrane and cytoplasm. Finally, the stabilization of endogenous of AMOT levels in p73KD EC treated with a tankyrase inhibitor [55] restored the junctional defects caused by p73-deficiency, altogether conclusively demonstrating that p73 regulates EC junction dynamics, at least in part, through the regulation of AMOT. This novel p73 role may not be exclusive of endothelial cells, since our data indicates that p73 regulates AMOT in different cell types, highlighting the need of addressing the relevance of the p73/AMOT/YAP regulatory axis in other cellular contexts in future investigations.

However, it is important to keep in mind that YAP and p73 functional interaction is complex. YAP interacts with p73 and behaves as its coactivator of pro-apoptotic genes [71]; in turn, p73 expression is essential for the recruitment of YAP to PML-nuclear bodies following DNA damage [71, 72], altogether suggesting that impaired p73 function could affect YAP activity at many levels in an AMOT-dependent and independent way. Thus, a deeper analysis would be required to fully untangle YAP/p73 functional interaction.

Knockdown of p73 in HUVEC directly impaired junctional and actin dynamics, affecting the establishment of both AJ and TJ, in particular reducing reticular junction formation. The idea of an interactive regulation between VE-Cadherin-mediated cell adhesion and actin dynamics in the control of barrier function has been previously introduced [67]. In the same line, reduction of reticular junctions affects VE-Cadherin remodeling [73] and has been linked to increased permeability in cultured EC [74]. Accordingly, p73 requirement for the establishment of VE-Cadherin reticular junctions could intertwine with TAp73 modulation of transcriptional programs regulating actin cytoskeleton dynamics [25] and would impact on endothelium integrity. Our observations not only reveal that p73 function is required for the formation of reticular junctions, but also to regulate the emergence of fingers, linking p73 function to the maintenance of the vascular barrier. Indeed, lack of p73 has functional consequences *in vitro* and *in vivo*, like enhanced permeability and hemorrhages, respectively. Therefore, p73 seems to play a predominant role in maintaining the integrity of the endothelial barrier in the absence of any stimuli. Endothelial barrier integrity is fundamental to ensure vascular and tissue homeostasis; consequently, in many diseases, the vascular barrier disintegrates, and leakage may become chronic. In cancer, blood vessels undergo changes fostered by the abnormal tumor microenvironment, including defective endothelial junctions that leads to increased permeability and leakage [75]. Based in our data it is feasible to speculate that p73 deregulation might play a role in the chronic vascular hyperpermeability associated with tumor angiogenesis [76] which represents a key point for future studies.

We propose a new model in which TAp73 acts as a vascular architect integrating transcriptional programs that will impinge with AMOT/YAP signaling to maintain junctional dynamics and integrity, whilst balancing endothelial cell rearrangements in angiogenic vessels. This situates p73 function in the hub of upstream regulators that control how endothelial cells establish and maintain adequate junctions to shape functional vascular networks, with important consequences in the maintenance of vascular homeostasis.

## Statement and Declarations

### Author Contributions

Conceptualization: L.M.-A., M.M.M. and M.C.M.; investigation: L.M.-A., H.A.-O., N.M.-G., J.V.-F., L.L.-F., L.P.-S., N.C.-A.; methodology: L.M.-A., J.V.-F., L.P.-S., J.M., L. d-P., L.C.-W., M.M.M. and M.C.M.; software: L.M.-A., H.A.-O., L.P.-S., validation: L.M.-A., J.M., L. d-P., M.M.M. and M.C.M.; formal analysis: L.M.-A., L. d-P., M.M.M. and M.C.M.; visualization: L.M.-A., H.A.-O., N.M.-G., M.M.M. and M.C.M.; writing the original draft, review and editing: L.M.-A., H.A.-O., M.M.M. and M.C.M.; general review and editing: All authors; supervision: L.M.-A., M.M.M. and M.C.M.; resources: A.F.-C., M.E.L.- M, B.J., L.H., M.W., J.M., L. d-P., L.C.-W., M.M.M. and M.C.M.; funding acquisition: M.C.M.; project administration: M.M.M. and M.C.M. All authors have read and agreed to the published version of the manuscript

## Data availability

The datasets generated during the current study are available in the OPEN SCAYLE repository https://open.scayle.es/en/dataset/maeso-alonso-et-al-2022

### Ethics approval

All animal experiments were carried out in agreement with European and Spanish regulations (Council Directive 2010/63/UE and RD 53/2013, respectively), with the appropriate institutional ethics committee approval (OEBA-ULE-018-2017 and OEBA-ULE-018-2021, Universidad de Leon and Junta de Castilla y Leon).

**Supplementary Figure 1.**
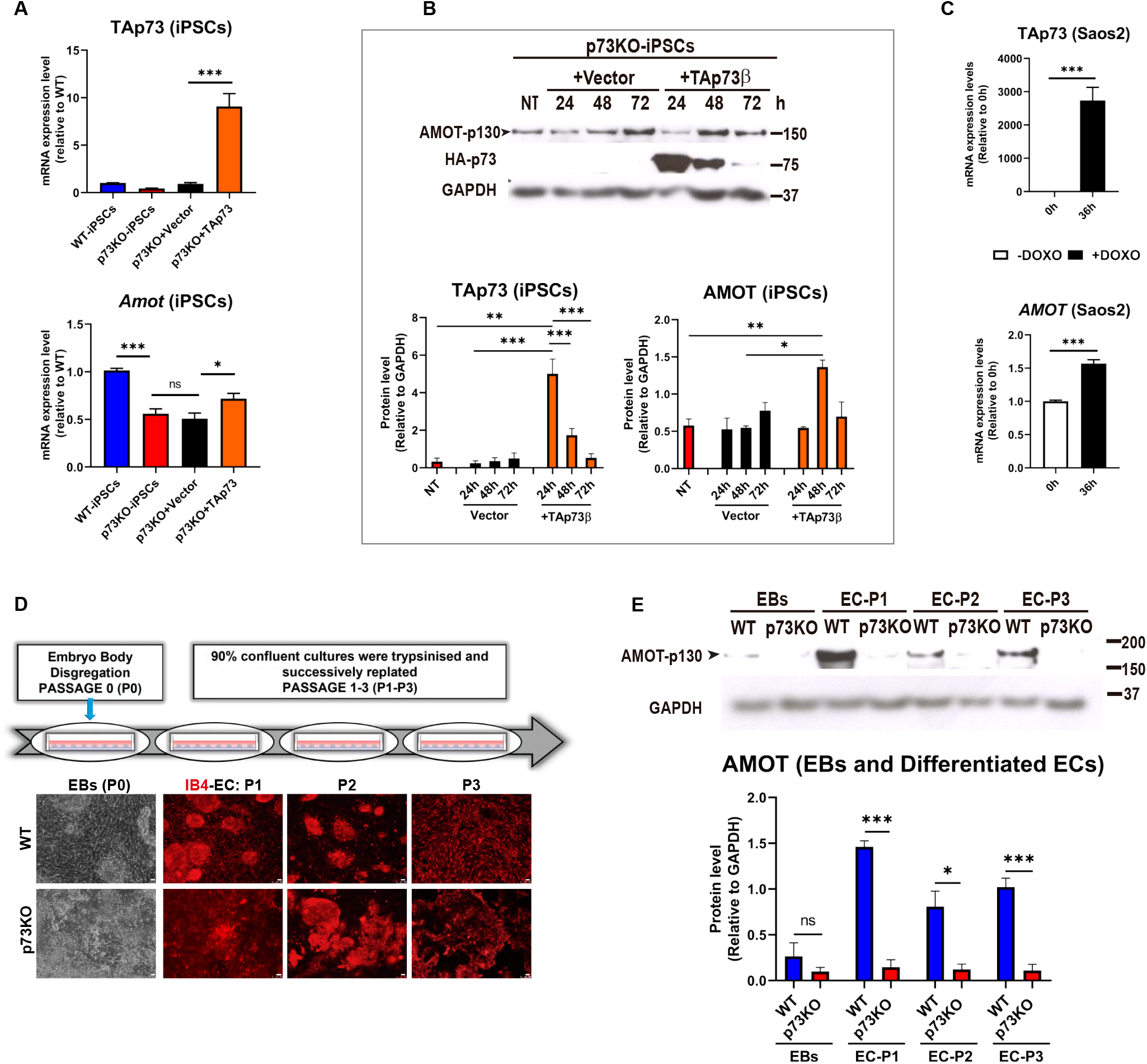
p73 regulates AMOT levels in different cell types. **(A, B)** Correlation of Amot and TAp73 expression in p73KO-iPSCs. Amot RNA (A) and protein (B) levels increased following TAp73 ectopic expression. For transfection experiments, iPSCs were seeded (3.7 × 10^4^ cells/cm^2^) on Matrigel™ (Corning, #356231) coated plates and transfected with an empty Vector or a TAp73 expression plasmid using Lipofectamine™ 3000 Transfection Reagent (Invitrogen, # L300008) following the manufacturer’s instructions. Cells were analysed 24 h after transfection (A) or up to 72 h after transfection (B). **(C)** TAp73 and *AMOT* expression analysis by qRT-PCR in HA-TAp73β-Saos2-Tet-On following Doxycycline treatment (–/+DOXO), which induces TAp73 expression. Bars in (A, B, C) represent the mean ± SEM of three independent experiments. *p<0.05, **p<0.01, ***p<0.001, ns: no significant. **(D)** Schematic drawing of the 2D-endothelial differentiation assay and representative phase-contrast and fluorescence microscopy images. EBs of the studied genotypes were disaggregated and plated (5 × 10^4^ cells/cm^2^) in EB medium (DMEM/GlutaMAX™-Gibco, #61965026, 25 mM HEPES, 1.2 mM sodium pyruvate, 19 mM monothioglycerol, 15% FBS) supplemented with 50 ng/ml VEGF-A (Peprotech, #450-32), on 0.1% gelatine-coated plates (non-endothelial, Passage 0). After 72-96 h hours in the presence of VEGF-A, 90% confluent cultures were trypsinised and replated on 0.1% gelatine-coated plates (endothelial differentiated cells, Passage 1). This step was repeated twice (Passage 2, 3). The cells were stained with biotinylated IB4 (2 μg/ml) to demonstrate their endothelial nature. Scale bar: 50 and 25 μm, respectively. **(E)** AMOT expression was analyzed by western blot in WT and p73KO undifferentiated cells (EBs) and quantified at various passages after endothelial differentiation (P1-P3).

**Supplementary Figure 2.**
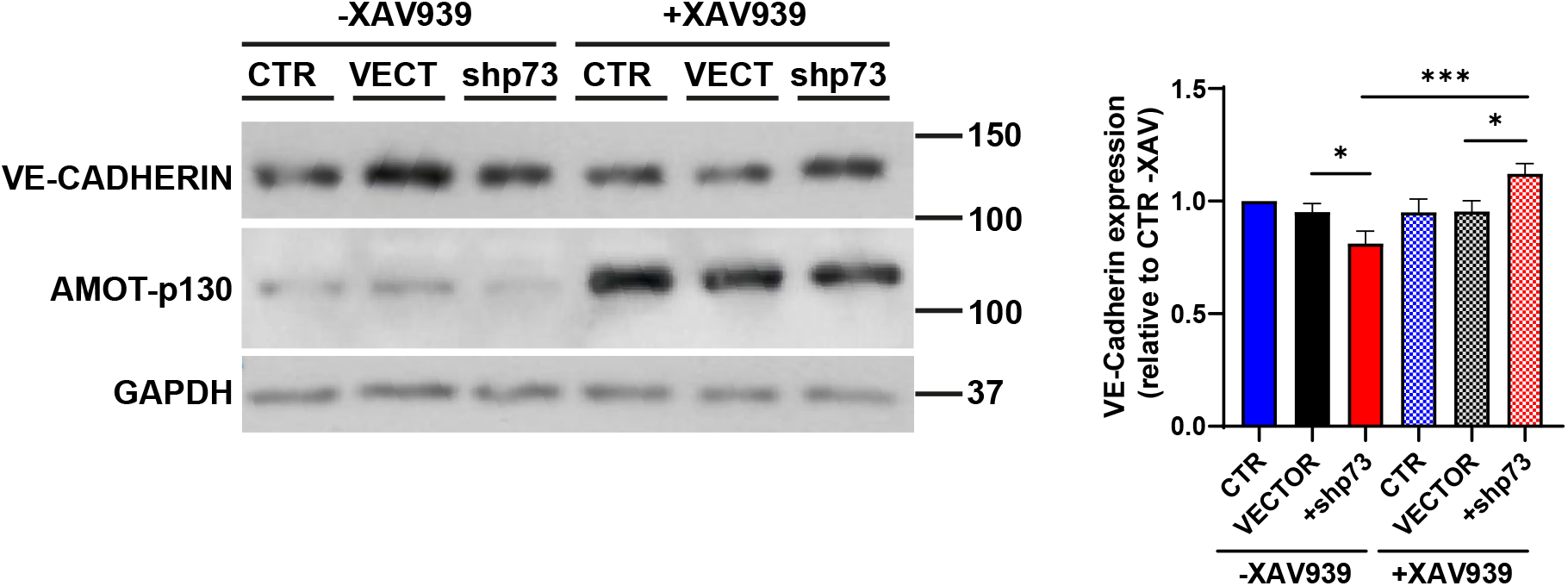
p73 knockdown has only a slight effect on VE-Cadherin expression. Western blot analysis of total VE-CADH in non-infected control HUVEC (CTR), and in cells infected with a non-silencing empty vector (VECT, VECTOR) or with an shRNA targeting p73 (+shp73). Following p73 knockdown (48 hours), cells were treated with XAV939 for 24 h. Treatment with the tankyrase inhibitor markedly induced AMOTp-130 expression. Only a small decrease of total VE-CADH expression was detected upon p73KD, although it was significantly recovered upon AMOT stabilization by XAV939 treatment. Graph shows the mean ± SEM of three independent experiments yielding similar results.

**Supplementary Figure 3.**
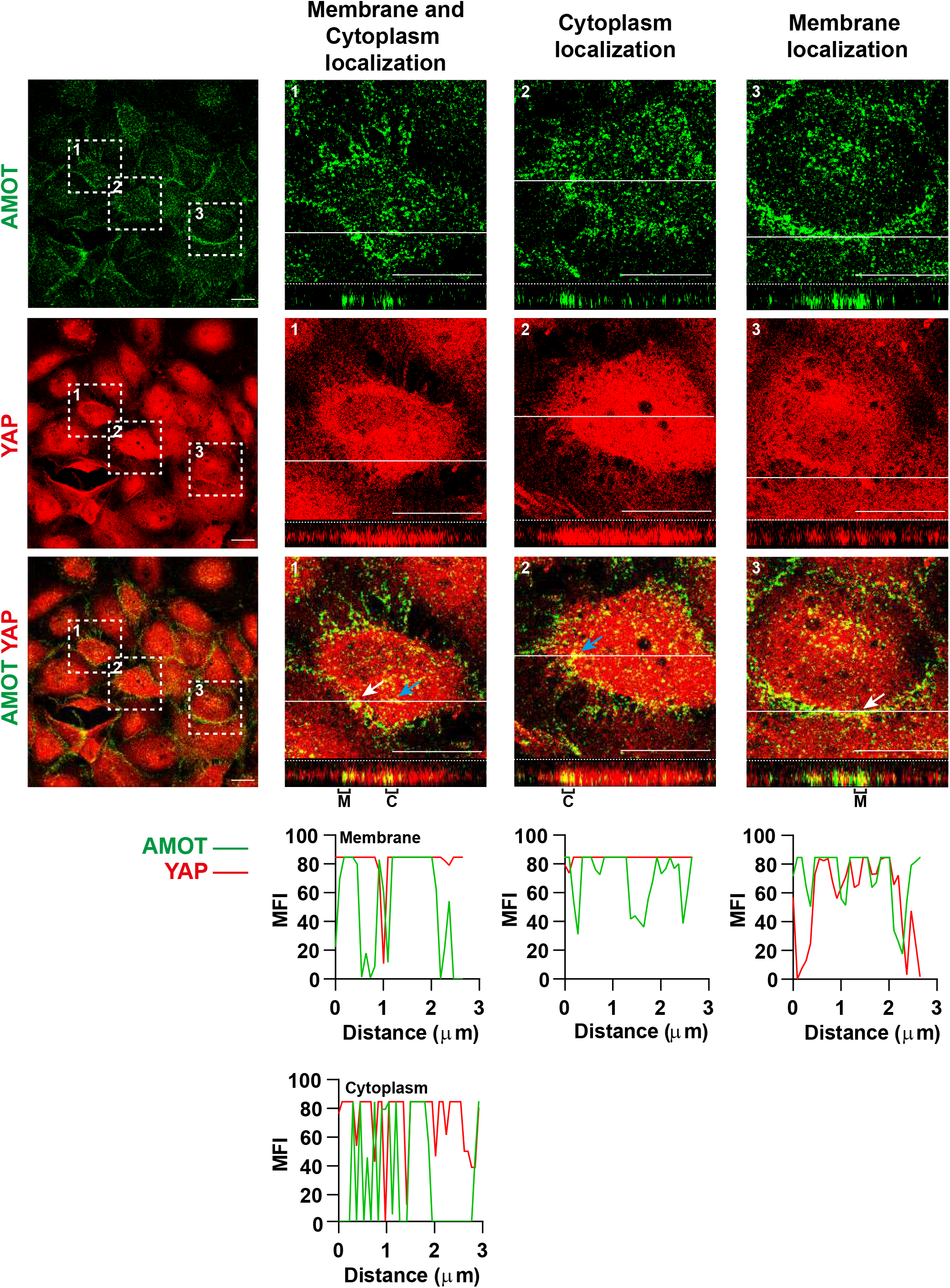
AMOT colocalizes with YAP at the plasma membrane and in the cytoplasm. Detailed analysis of selected areas (white dashed squares 1, 2, 3) from Figure 5 confocal micrographs showing confluent monolayers of p73KD cells treated with the tankyrase inhibitor XAV939. Cells were immunostained for AMOT (green) and YAP (red). Orthogonal projections of the magnified areas (1, 2, 3) show AMOT staining at the plasma membrane (M bracket) or the cytoplasm (C bracket) colocalizing with YAP. The histograms represent the mean fluorescence intensity (MFI) profiles of AMOT and YAP expression in the regions indicated by brackets at the orthogonal projections. Scale bar: 20 µm.

**Supplementary Figure 4.**
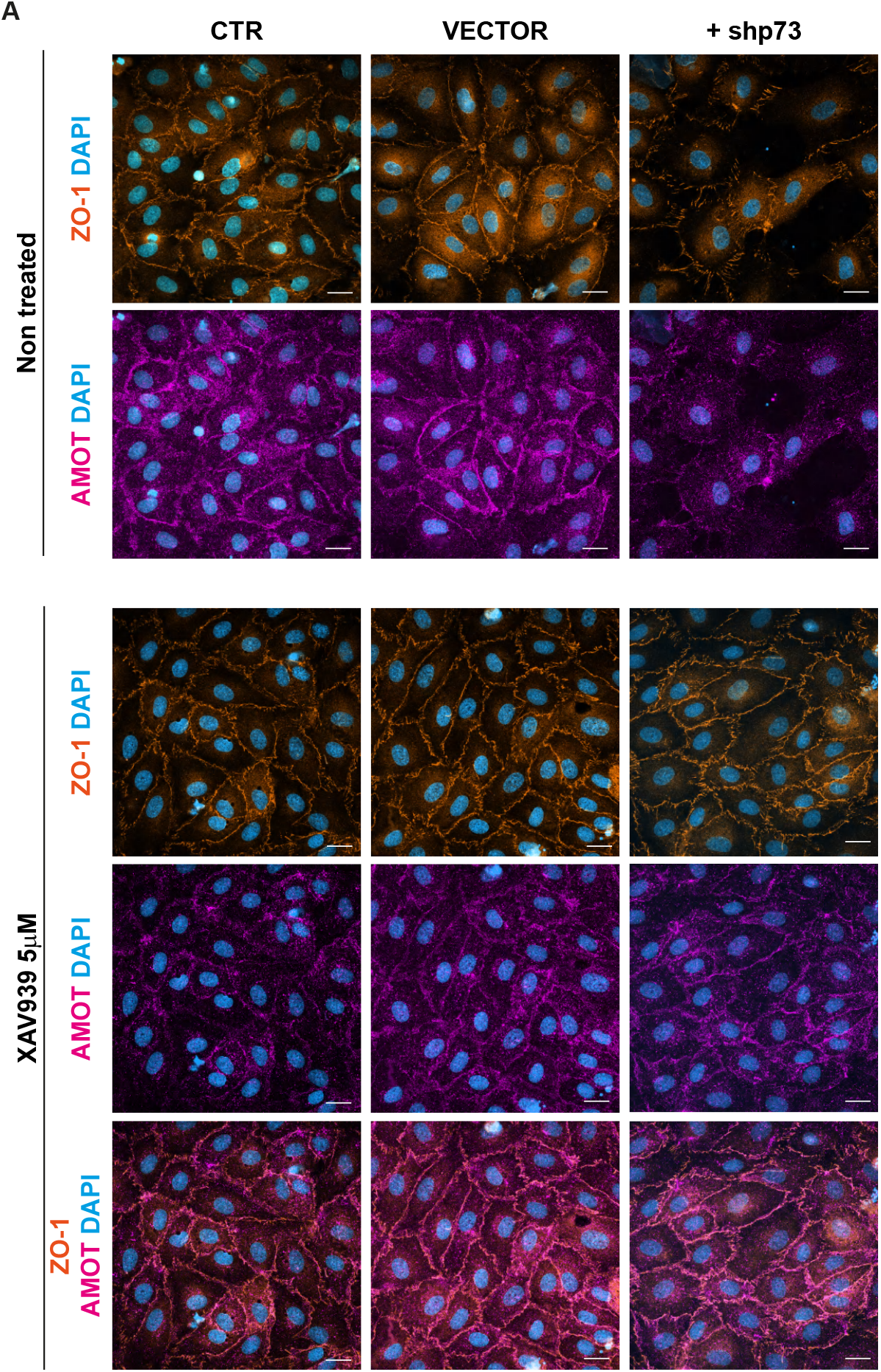
AMOT is a p73-downstream effector for the regulation of tight junctions. Stabilization of endogenous AMOT levels by XAV939 treatment restores the tight junctions defects induced by the p73KD. AMOT (pink) restoration rescues the morphology of TJs as shown by ZO-1 (orange) staining. Nuclei were counterstained with DAPI (blue). Scale bar: 20 µm.

