## Supplementary material for "p73 is required for vessel integrity controlling endothelial junctional dynamics through Angiomotin": Supp Table 1

**Supplementary Table 1. List of DEGs identified by RNA-seq in p73KO-iPSCs after TAp73β ectopic expression.** DEGs were consider significant at p-adj<0.05

| Gene | log2FC | p-adj |
|---|---|---|
| *Krt14* | 2,69 | 9,03E-96 |
| *Foxj1* | 2,55 | 1,45E-92 |
| *Perp* | 1,68 | 4,49E-71 |
| *Cdkn1c* | 1,42 | 1,66E-34 |
| *Cemip* | 1,59 | 3,63E-32 |
| *Itgb4* | 1,45 | 4,86E-32 |
| *Mdm2* | 0,91 | 1,91E-31 |
| *Dst* | 0,83 | 9,87E-22 |
| *Cbr2* | 1,09 | 1,27E-17 |
| *Junb* | 0,93 | 8,71E-17 |
| *Bcam* | 0,62 | 2,74E-16 |
| *Sord* | 1,01 | 1,24E-15 |
| *Ngfr* | 0,92 | 1,64E-14 |
| *Trp73* | 1,04 | 2,04E-14 |
| *Nptxr* | 1,05 | 5,21E-14 |
| *Gata6* | 0,96 | 2,44E-13 |
| *Idh1* | 0,69 | 3,38E-13 |
| *Pak6* | 1,00 | 1,10E-12 |
| *Wnt9a* | 0,94 | 3,01E-12 |
| *Vwa1* | 0,96 | 6,39E-12 |
| *Anxa8* | 0,97 | 7,04E-12 |
| *Jag2* | 0,91 | 1,03E-11 |
| *Ass1* | 0,49 | 1,98E-11 |
| *Cd109* | 0,92 | 2,78E-11 |
| *Itga6* | 0,54 | 3,59E-11 |
| *Wnt4* | 0,88 | 9,55E-11 |
| *Krt17* | 0,91 | 1,22E-10 |
| *Fermt1* | 0,93 | 1,31E-10 |
| *Cx3cl1* | 0,93 | 1,80E-10 |
| *Cotl1* | 0,76 | 2,57E-10 |
| *Slc38a3* | 0,87 | 4,19E-10 |
| *Amotl1* | 0,69 | 1,45E-09 |
| *Mfge8* | 0,58 | 2,15E-09 |
| *Scube3* | 0,81 | 2,15E-09 |
| *Col18a1* | 0,42 | 8,79E-09 |
| *Erf* | 0,48 | 8,88E-09 |
| *Lama5* | 0,54 | 1,18E-08 |
| *Cic* | 0,49 | 4,21E-08 |
| *Pard6g* | 0,81 | 6,22E-08 |
| *Lars2* | 0,36 | 8,10E-08 |
| *Apoe* | 0,43 | 8,14E-08 |
| *Gm26917* | 0,46 | 1,15E-07 |
| *Fermt3* | 0,70 | 1,95E-07 |
| *Tjp3* | 0,77 | 4,55E-07 |
| *Capn1* | 0,65 | 4,86E-07 |
| *Pou4f3* | 0,65 | 4,86E-07 |
| *Card10* | 0,69 | 6,49E-07 |
| *Cacna2d2* | 0,70 | 6,51E-07 |
| *Sytl1* | 0,74 | 6,51E-07 |
| *Col7a1* | 0,75 | 1,18E-06 |

| Gene | log2FC | p-adj |
|---|---|---|
| *Txnip* | -0,40 | 2,12E-06 |
| *Fam101b* | 0,71 | 2,46E-06 |
| *Galnt18* | 0,67 | 2,88E-06 |
| *Coro6* | 0,59 | 4,19E-06 |
| *Tgfbr1* | 0,61 | 5,49E-06 |
| *Bmp7* | 0,68 | 5,71E-06 |
| *Zfp42* | -0,35 | 6,14E-06 |
| *Plch2* | 0,63 | 9,10E-06 |
| *Bbx* | 0,56 | 9,13E-06 |
| *Fgf4* | -0,41 | 9,13E-06 |
| *Fxyd3* | 0,51 | 9,96E-06 |
| *Nid2* | 0,39 | 9,98E-06 |
| *Nanog* | -0,35 | 1,09E-05 |
| *Gas6* | 0,69 | 1,19E-05 |
| *Exoc3l4* | 0,62 | 1,23E-05 |
| *Fam129b* | 0,40 | 1,23E-05 |
| *Phc1* | -0,32 | 1,31E-05 |
| *Rapgefl1* | 0,65 | 1,39E-05 |
| *Tnrc18* | 0,55 | 1,67E-05 |
| *Krtdap* | 0,60 | 2,00E-05 |
| *Krt19* | 0,67 | 2,82E-05 |
| *Pdgfa* | 0,57 | 2,82E-05 |
| *Tdgf1* | -0,31 | 3,71E-05 |
| *Lamb1* | 0,46 | 4,00E-05 |
| *Ldlrap1* | 0,62 | 4,31E-05 |
| *Mcam* | -0,48 | 4,31E-05 |
| *Mbp* | 0,64 | 4,51E-05 |
| *Ivl* | 0,46 | 5,15E-05 |
| *Nr0b1* | -0,44 | 5,21E-05 |
| *Fam83g* | 0,65 | 6,19E-05 |
| *Tagln2* | 0,38 | 8,23E-05 |
| *Frmd4b* | 0,64 | 9,51E-05 |
| *Rdh10* | 0,64 | 9,51E-05 |
| *Spp1* | -0,31 | 9,51E-05 |
| *Hr* | 0,63 | 1,07E-04 |
| *Ptpn13* | 0,57 | 1,07E-04 |
| *Wnt7b* | 0,54 | 1,17E-04 |
| *Esrrb* | -0,29 | 1,20E-04 |
| *Spon2* | 0,49 | 1,36E-04 |
| *Tex261* | 0,49 | 1,36E-04 |
| *Aldh1a3* | 0,61 | 1,37E-04 |
| *Vgf* | 0,62 | 1,37E-04 |
| *mt-Atp8* | 0,44 | 1,39E-04 |
| *Eef2k* | 0,59 | 1,54E-04 |
| *Dnmt3a* | -0,35 | 1,65E-04 |
| *Irgm1* | -0,44 | 1,68E-04 |
| *Sdc4* | 0,38 | 1,97E-04 |
| *Mir6236* | 0,48 | 2,00E-04 |
| *Gm15662* | 0,42 | 2,06E-04 |
| *Zfp57* | -0,33 | 2,06E-04 |

| Gene | log2FC | padj |
|---|---|---|
| *Rnf144b* | 0,68 | 1,18E-06 |
| *Kremen1* | 0,68 | 1,33E-06 |
| *Jarid2* | -0,33 | 1,41E-06 |
| *Gprc5c* | 0,74 | 1,46E-06 |
| *Limk2* | 0,48 | 1,85E-06 |
| *Eps8l2* | 0,59 | 3,26E-04 |
| *Ece1* | 0,44 | 3,35E-04 |
| *Abhd17c* | 0,46 | 3,36E-04 |
| *Wnt3a* | 0,51 | 3,42E-04 |
| *Mkrn1* | -0,26 | 3,46E-04 |
| *Adam8* | 0,59 | 3,56E-04 |
| *Fam83f* | 0,47 | 3,57E-04 |
| *Nbeal2* | 0,57 | 3,78E-04 |
| *Pkp3* | 0,58 | 3,78E-04 |
| *Dusp2* | 0,46 | 4,11E-04 |
| *Tmppe* | 0,52 | 4,29E-04 |
| *Il1rl2* | 0,45 | 4,33E-04 |
| *Coro7* | 0,58 | 4,50E-04 |
| *Sncg* | 0,53 | 4,91E-04 |
| *Pltp* | 0,55 | 4,97E-04 |
| *Lamb3* | 0,55 | 4,98E-04 |
| *RP23-473E2(* | 0,53 | 6,51E-04 |
| *Slc12a4* | 0,44 | 6,61E-04 |
| *Fat2* | 0,45 | 7,53E-04 |
| *Sertad4* | 0,52 | 9,01E-04 |
| *Glb1* | 0,48 | 9,43E-04 |
| *Pim1* | -0,38 | 9,64E-04 |
| *Cpa4* | 0,37 | 1,07E-03 |
| *Pdia6* | 0,27 | 1,25E-03 |
| *Dmkn* | 0,54 | 1,28E-03 |
| *Vwa2* | 0,55 | 1,29E-03 |
| *Sept1* | -0,33 | 1,43E-03 |
| *Bcl9l* | 0,54 | 1,47E-03 |
| *Oasl2* | -0,55 | 1,63E-03 |
| *Ddx58* | -0,36 | 1,71E-03 |
| *Hagh* | 0,54 | 1,71E-03 |
| *Isg15* | -0,53 | 1,71E-03 |
| *Myrf* | -0,43 | 1,71E-03 |
| *Hspg2* | 0,34 | 1,77E-03 |
| *Ifit1* | -0,55 | 1,77E-03 |
| *Lima1* | 0,47 | 2,03E-03 |
| *Lamc2* | 0,45 | 2,06E-03 |
| *Rcan1* | 0,50 | 2,17E-03 |
| *Bend4* | 0,41 | 2,46E-03 |
| *Spry4* | -0,38 | 2,52E-03 |
| *Mark4* | 0,48 | 2,68E-03 |
| *Chst3* | 0,47 | 2,74E-03 |
| *Krt5* | 0,37 | 2,78E-03 |
| *Ptp4a3* | -0,45 | 2,78E-03 |
| *Cebpa* | 0,44 | 2,90E-03 |
| *Elmsan1* | 0,44 | 3,03E-03 |
| *Mvb12b* | 0,52 | 3,03E-03 |
| *Ubald2* | 0,34 | 3,08E-03 |
| *Mafb* | 0,40 | 3,13E-03 |
| *Parp12* | -0,44 | 3,16E-03 |

| Gene | log2FC | padj |
|---|---|---|
| *Sema3f* | 0,57 | 2,10E-04 |
| *Pxn* | 0,47 | 2,90E-04 |
| *Sept9* | 0,37 | 3,00E-04 |
| *Dusp7* | 0,59 | 3,07E-04 |
| *Csrp1* | 0,38 | 3,23E-04 |
| *Efs* | 0,46 | 3,94E-03 |
| *Cmip* | 0,36 | 3,99E-03 |
| *Slc7a1* | 0,34 | 4,30E-03 |
| *Phyhip* | 0,42 | 4,38E-03 |
| *Dppa5a* | -0,21 | 4,68E-03 |
| *L1td1* | -0,21 | 4,68E-03 |
| *P2ry1* | 0,37 | 4,68E-03 |
| *Dab2ip* | 0,37 | 4,70E-03 |
| *Mt2* | -0,25 | 4,80E-03 |
| *Myh9* | 0,27 | 4,80E-03 |
| *Ngfrap1* | 0,29 | 4,80E-03 |
| *Marcksl1* | 0,26 | 4,84E-03 |
| *Nxn* | 0,43 | 4,87E-03 |
| *Jup* | 0,29 | 5,55E-03 |
| *Glul* | -0,30 | 5,68E-03 |
| *Plxnb2* | 0,33 | 5,77E-03 |
| *Efnb1* | 0,48 | 6,37E-03 |
| *Slc29a1* | -0,29 | 6,52E-03 |
| *Itpkb* | 0,47 | 6,57E-03 |
| *Parp9* | -0,46 | 6,59E-03 |
| *AI661453* | 0,46 | 7,20E-03 |
| *Gjb3* | 0,35 | 7,42E-03 |
| *Dag1* | 0,26 | 8,34E-03 |
| *Cyp26b1* | 0,40 | 8,48E-03 |
| *Rgs16* | -0,48 | 8,48E-03 |
| *Fads2* | 0,43 | 8,72E-03 |
| *Pcsk5* | 0,32 | 9,00E-03 |
| *Mmp9* | 0,43 | 9,13E-03 |
| *Rbpms2* | -0,31 | 9,18E-03 |
| *Atp13a3* | 0,27 | 9,20E-03 |
| *Sall1* | -0,34 | 9,23E-03 |
| *Nav2* | 0,33 | 9,36E-03 |
| *Slc7a3* | -0,32 | 9,42E-03 |
| *Trim47* | 0,45 | 9,58E-03 |
| *Il6ra* | 0,45 | 9,59E-03 |
| *Hyou1* | 0,26 | 9,81E-03 |
| *Irf6* | 0,48 | 9,81E-03 |
| *Hes2* | 0,34 | 9,95E-03 |
| *Stard8* | 0,49 | 9,95E-03 |
| *Pou5f1* | -0,22 | 9,98E-03 |
| *Grem1* | -0,38 | 0,010 |
| *Ppp1r13l* | 0,48 | 0,010 |
| *Cd151* | 0,36 | 0,010 |
| *Pvrl1* | 0,46 | 0,010 |
| *Wnt11* | 0,35 | 0,011 |
| *Nasp* | -0,20 | 0,011 |
| *Phldb1* | 0,45 | 0,012 |
| *Sfn* | 0,42 | 0,012 |
| *Hyal2* | 0,42 | 0,012 |
| *Ttc7b* | 0,42 | 0,012 |

| Gene | log2FC | padj |
|---|---|---|
| *Rarg* | -0,30 | 3,25E-03 |
| *Cd276* | 0,47 | 3,56E-03 |
| *Ccnd1* | -0,34 | 3,66E-03 |
| *Fam60a* | -0,27 | 3,74E-03 |
| *Rnase4* | 0,45 | 3,74E-03 |
| *Nsmaf* | 0,43 | 0,014 |
| *Tmem94* | 0,44 | 0,014 |
| *Fam83h* | 0,42 | 0,014 |
| *Hip1r* | 0,47 | 0,014 |
| *Rgs3* | 0,44 | 0,015 |
| *Alox12* | 0,35 | 0,016 |
| *Dsp* | 0,39 | 0,016 |
| *Hmgxb4* | -0,27 | 0,016 |
| *Oas2* | -0,46 | 0,018 |
| *Itga3* | 0,46 | 0,019 |
| *Bmp6* | 0,36 | 0,019 |
| *Flnb* | 0,28 | 0,019 |
| *Enah* | -0,21 | 0,019 |
| *Etv5* | -0,23 | 0,020 |
| *2900026A02R* | 0,46 | 0,020 |
| *Dhrs3* | 0,37 | 0,020 |
| *Src* | 0,36 | 0,020 |
| *Dhx16* | -0,25 | 0,021 |
| *Slc25a4* | 0,29 | 0,021 |
| *Ackr3* | 0,45 | 0,022 |
| *Ctdspl* | 0,43 | 0,023 |
| *Id1* | 0,40 | 0,023 |
| *Krt18* | 0,43 | 0,023 |
| *Gltscr1l* | -0,32 | 0,024 |
| *Tgfa* | 0,36 | 0,024 |
| *Sh3pxd2a* | 0,37 | 0,024 |
| *Tdh* | -0,25 | 0,024 |
| *Lsp1* | 0,39 | 0,024 |
| *Trim6* | -0,24 | 0,025 |
| *Hsp90b1* | 0,19 | 0,026 |
| *Hmcn2* | 0,43 | 0,026 |
| *Pml* | -0,27 | 0,026 |
| *Fem1b* | 0,23 | 0,026 |
| *Micall1* | 0,44 | 0,028 |
| *Esrp2* | 0,44 | 0,028 |
| *Sox2* | -0,27 | 0,028 |
| *Gadd45a* | -0,36 | 0,028 |
| *Capg* | 0,38 | 0,028 |
| *Sema4b* | 0,29 | 0,028 |
| *Dlx3* | 0,34 | 0,029 |
| *Mapkbp1* | 0,44 | 0,029 |
| *Myo1c* | 0,27 | 0,029 |
| *Gm42927* | -0,22 | 0,029 |
| *Vim* | -0,26 | 0,029 |
| *Gfpt2* | -0,32 | 0,029 |
| *Kctd12* | 0,40 | 0,029 |
| *Hspa5* | 0,19 | 0,030 |
| *Rcor2* | -0,28 | 0,030 |
| *Tacstd2* | 0,43 | 0,031 |
| *Hras* | 0,35 | 0,032 |

| Gene | log2FC | padj |
|---|---|---|
| *Tet1* | -0,21 | 0,032 |
| *Trp53inp1* | -0,27 | 0,033 |
| *Zfp36l2* | 0,32 | 0,033 |
| *1300017J02R* | 0,32 | 0,035 |
| *Lrrn4* | 0,43 | 0,035 |
| *Scn4b* | 0,33 | 0,012 |
| *Cnn2* | 0,39 | 0,013 |
| *Vegfa* | 0,48 | 0,013 |
| *Myo18a* | 0,37 | 0,013 |
| *Helz2* | -0,47 | 0,014 |
| *Tns4* | 0,42 | 0,035 |
| *Mbnl2* | 0,36 | 0,035 |
| *Epha1* | 0,43 | 0,036 |
| *Sema4c* | 0,39 | 0,036 |
| *Sh3bgrl2* | 0,37 | 0,036 |
| *Exoc3l2* | 0,33 | 0,036 |
| *Rev1* | -0,33 | 0,036 |
| *Setd7* | 0,38 | 0,037 |
| *Plek2* | 0,40 | 0,037 |
| *Trim25* | -0,22 | 0,038 |
| *Nrg1* | 0,43 | 0,038 |
| *Pdpn* | 0,35 | 0,038 |
| *Hs6st1* | 0,28 | 0,040 |
| *Tulp4* | 0,40 | 0,042 |
| *Enc1* | -0,23 | 0,042 |
| *Fam89a* | 0,30 | 0,042 |
| *Dgka* | 0,35 | 0,044 |
| *Mcm10* | -0,27 | 0,044 |
| *S100a11* | 0,33 | 0,046 |
| *Acp6* | -0,34 | 0,047 |
| *Edaradd* | 0,24 | 0,048 |
| *Rgs14* | 0,32 | 0,048 |
| *Ifrd1* | -0,26 | 0,048 |
| *Adamts1* | -0,40 | 0,049 |
| *Calr* | 0,19 | 0,0496 |
| *Duox1* | 0,25 | 0,0496 |
