## Supplementary material for "p73 is required for vessel integrity controlling endothelial junctional dynamics through Angiomotin": Supp Table 2

**Supplementary Table 2. List of significant Biological Process (FDR<0.05) in p73KO-iPSCs after TAp73β ectopic expression.** FE: Fold Enrichment. FDR: Fold Discovery Rate

| Term | Count | Genes | FE | FDR |
|---|---|---|---|---|
| GO:0007275~multicellular organism development | 115 | PDGFA, MMP9, WNT3A, BBX, JAG2, ITPKB, WNT4, GATA6, APOE, ADAM8, EDARADD, DAB2IP, PDPN, PLXNB2, EFNB1, MFGE8, MYH9, MARK4, JUNB, JUP, KRT19, KRT17, TACSTD2, NAV2, VEGFA, DSP, LAMC2, TRP73, DST, SLC38A3, ITGB4, DAG1, CX3CL1, CALR, PXN, SRC, LAMB3, ECE1, P2RY1, IDH1, LAMB1, IVL, COL18A1, MAFB, TGFBR1, IL1RL2, CD276, ITGA3, EPHA1, GAS6, CDKN1C, DLX3, NXN, ITGA6, LAMA5, BMP7, BMP6, HIP1R, DUOX1, SDC4, VGF, MBP, SEMA3F, POU4F3, EEF2K, TGFA, FAM83H, NRG1, HYAL2, SH3PXD2A, SLC25A4, FOXJ1, ACKR3, FLNB, DHRS3, NPTXR, SEMA4C, MDM2, SEMA4B, WNT11, ESRP2, NGFR, WNT9A, ALOX12, PHLDB1, CD109, SFN, FEM1B, CIC, ZFP36L2, COL7A1, ALDH1A3, CYP26B1, BCL9L, CNN2, HS6ST1, HSPA5, NBEAL2, PCSK5, CEBPA, ERF, GJB3, HSPG2, RCAN1, PPP1R13L, RGS14, MICALL1, IL6RA, CAPN1, WNT7B, RDH10, DUSP2, IRF6, ID1, PERP | 1,88 | 1,80E-11 |
| GO:0048731~system development | 109 | PDGFA, MMP9, WNT3A, BBX, JAG2, ITPKB, WNT4, GATA6, APOE, ADAM8, EDARADD, DAB2IP, PDPN, PLXNB2, EFNB1, MFGE8, MYH9, MARK4, JUNB, JUP, KRT19, KRT17, TACSTD2, NAV2, VEGFA, DSP, LAMC2, TRP73, DST, SLC38A3, ITGB4, DAG1, CX3CL1, CALR, SRC, ECE1, P2RY1, IDH1, LAMB1, IVL, COL18A1, MAFB, IL1RL2, TGFBR1, CD276, ITGA3, EPHA1, GAS6, CDKN1C, DLX3, ITGA6, NXN, LAMA5, BMP7, BMP6, HIP1R, SDC4, VGF, MBP, SEMA3F, POU4F3, EEF2K, TGFA, FAM83H, NRG1, HYAL2, SH3PXD2A, SLC25A4, FOXJ1, ACKR3, FLNB, DHRS3, NPTXR, SEMA4C, MDM2, SEMA4B, WNT11, NGFR, WNT9A, ESRP2, ALOX12, CD109, SFN, FEM1B, CIC, ZFP36L2, ALDH1A3, CYP26B1, BCL9L, CNN2, HS6ST1, HSPA5, NBEAL2, PCSK5, CEBPA, ERF, GJB3, HSPG2, RCAN1, PPP1R13L, RGS14, CAPN1, IL6RA, MICALL1, WNT7B, RDH10, IRF6, ID1, PERP | 2,01 | 1,72E-12 |

| | | | | |
|---|---|---|---|---|
| GO:0048869~cellular developmental process | 96 | HRAS, FERMT3, MMP9, WNT3A, JAG2, ITPKB, SDC4, WNT4, GATA6, APOE, SEMA3F, DMKN, POU4F3, EEF2K, ADAM8, NRG1, EDARADD, HYAL2, DAB2IP, SH3PXD2A, SLC25A4, PDPN, FOXJ1, EFNB1, PLXNB2, KRTDAP, ACKR3, MYH9, FLNB, JUNB, KRT19, NPTXR, KRT17, TACSTD2, KRT14, VEGFA, SEMA4C, MDM2, SEMA4B, DSP, WNT11, TRP73, NGFR, WNT9A, DST, SEPT9, ALOX12, PHLDB1, ITGB4, DAG1, CD109, CX3CL1, SFN, CALR, TAGLN2, FEM1B, PXN, SRC, ZFP36L2, LAMB3, COL7A1, CYP26B1, P2RY1, BCL9L, HS6ST1, HSPA5, LAMB1, NBEAL2, IVL, COL18A1, CEBPA, ERF, MAFB, TGFBR1, IL1RL2, HSPG2, CD276, RCAN1, ITGA3, PPP1R13L, CACNA2D2, EPHA1, RGS14, GAS6, MICALL1, DLX3, CDKN1C, WNT7B, RDH10, NXN, ITGA6, IRF6, ID1, LAMA5, BMP7, BMP6 | 1,83 | 1,56E-07 |
| GO:0048513~animal organ development | 92 | PDGFA, MMP9, WNT3A, BBX, JAG2, ITPKB, SDC4, VGF, MBP, WNT4, GATA6, SEMA3F, POU4F3, TGFA, ADAM8, FAM83H, NRG1, EDARADD, HYAL2, DAB2IP, SH3PXD2A, SLC25A4, PDPN, FOXJ1, EFNB1, PLXNB2, MYH9, FLNB, JUNB, JUP, DHRS3, KRT19, KRT17, TACSTD2, VEGFA, SEMA4C, MDM2, SEMA4B, DSP, WNT11, TRP73, NGFR, ESRP2, WNT9A, ALOX12, SLC38A3, ITGB4, DAG1, CD109, SFN, CALR, FEM1B, CIC, SRC, ZFP36L2, ECE1, ALDH1A3, CYP26B1, IDH1, BCL9L, CNN2, HS6ST1, HSPA5, LAMB1, PCSK5, NBEAL2, IVL, CEBPA, ERF, MAFB, TGFBR1, IL1RL2, GJB3, HSPG2, CD276, ITGA3, RCAN1, PPP1R13L, GAS6, CAPN1, IL6RA, DLX3, CDKN1C, WNT7B, RDH10, ITGA6, IRF6, LAMA5, ID1, PERP, BMP7, BMP6 | 2,22 | 4,98E-12 |
| GO:0009653~anatomical structure morphogenesis | 85 | PDGFA, FERMT3, MMP9, WNT3A, JAG2, SDC4, WNT4, GATA6, APOE, SEMA3F, POU4F3, EEF2K, TGFA, ADAM8, NRG1, EDARADD, HYAL2, DAB2IP, SH3PXD2A, PDPN, FOXJ1, EFNB1, PLXNB2, ACKR3, MFGE8, MYH9, FLNB, JUNB, KRT19, DHRS3, KRT17, TACSTD2, VEGFA, SEMA4C, MDM2, SEMA4B, DSP, WNT11, TRP73, NGFR, ESRP2, WNT9A, DST, SEPT9, ALOX12, PHLDB1, ITGB4, DAG1, CD109, CX3CL1, CALR, FEM1B, SRC, PXN, LAMB3, ECE1, COL7A1, ALDH1A3, CYP26B1, BCL9L, HS6ST1, HSPA5, LAMB1, PCSK5, NBEAL2, COL18A1, ERF, MAFB, TGFBR1, HSPG2, ITGA3, PPP1R13L, EPHA1, CAPN1, DLX3, CDKN1C, RDH10, WNT7B, DUSP2, ITGA6, LAMA5, ID1, PERP, BMP7, BMP6 | 2,63 | 3,36E-15 |

| | | | | |
|---|---|---|---|---|
| GO:0009888~tissue development | 73 | PDGFA, WNT3A, MMP9, JAG2, SDC4, WNT4, GATA6, SEMA3F, POU4F3, ADAM8, FAM83H, NRG1, EDARADD, HYAL2, DAB2IP, SLC25A4, PDPN, FOXJ1, EFNB1, PLXNB2, KRTDAP, FLNB, JUNB, KRT17, TACSTD2, VEGFA, KRT14, SEMA4C, MDM2, SEMA4B, DSP, WNT11, TRP73, NGFR, WNT9A, ESRP2, ALOX12, ITGB4, CD109, DAG1, SFN, CALR, TAGLN2, FEM1B, SRC, PXN, LAMB3, COL7A1, ALDH1A3, CYP26B1, BCL9L, LAMB1, IVL, COL18A1, ERF, TGFBR1, HSPG2, CD276, ITGA3, RCAN1, PPP1R13L, GAS6, DLX3, CDKN1C, RDH10, WNT7B, DUSP2, ITGA6, IRF6, LAMA5, ID1, BMP7, BMP6 | 3,28 | 1,77E-17 |
| GO:0010646~regulation of cell communication | 66 | SNCG, HRAS, PDGFA, HIP1R, WNT3A, MMP9, JAG2, ITPKB, VGF, WNT4, GATA6, APOE, TGFA, ADAM8, NRG1, MAPKBP1, VWA2, MVB12B, HYAL2, DAB2IP, SCUBE3, ACKR3, CARD10, DHRS3, VEGFA, SEMA4C, MDM2, WNT11, TRP73, NGFR, WNT9A, EPS8L2, CD109, DAG1, CX3CL1, CALR, FEM1B, SRC, PXN, ECE1, ALDH1A3, CYP26B1, P2RY1, BCL9L, HSPA5, MYO1C, TGFBR1, PTPN13, ITGA3, RCAN1, CACNA2D2, RGS14, GAS6, IL6RA, CDKN1C, HYOU1, WNT7B, ITGA6, NXN, ID1, RGS3, SLC7A1, BMP7, DUSP7, KCTD12, BMP6 | 2,01 | 2,14E-05 |
| GO:0009966~regulation of signal transduction | 63 | HRAS, PDGFA, HIP1R, WNT3A, MMP9, JAG2, ITPKB, WNT4, GATA6, APOE, TGFA, ADAM8, NRG1, MAPKBP1, VWA2, MVB12B, DAB2IP, HYAL2, SCUBE3, ACKR3, CARD10, DHRS3, VEGFA, SEMA4C, MDM2, WNT11, TRP73, NGFR, WNT9A, EPS8L2, CD109, DAG1, CX3CL1, CALR, FEM1B, SRC, PXN, ECE1, ALDH1A3, CYP26B1, P2RY1, BCL9L, HSPA5, MYO1C, TGFBR1, PTPN13, ITGA3, RCAN1, RGS14, GAS6, IL6RA, CDKN1C, HYOU1, WNT7B, ITGA6, NXN, ID1, RGS3, SLC7A1, BMP7, DUSP7, KCTD12, BMP6 | 2,17 | 3,14E-06 |
| GO:0006928~movement of cell or subcellular component | 62 | HRAS, PDGFA, FERMT3, WNT3A, MMP9, FERMT1, AMOTL1, SDC4, CD151, WNT4, APOE, SEMA3F, POU4F3, ADAM8, FAM83H, NRG1, DAB2IP, HYAL2, PDPN, FOXJ1, EFNB1, PLXNB2, CHST3, MYH9, JUP, TACSTD2, VEGFA, CEMIP, SEMA4C, MDM2, SEMA4B, DSP, LAMC2, WNT11, NGFR, DST, MYO18A, ALOX12, SORD, DAG1, ITGB4, CX3CL1, CALR, SRC, PXN, FAT2, P2RY1, CNN2, HSPA5, LAMB1, PLTP, COL18A1, MYO1C, TGFBR1, ITGA3, EPHA1, GAS6, WNT7B, ITGA6, LAMA5, SCN4B, BMP7 | 3,05 | 1,89E-12 |

| | | | | |
|---|---|---|---|---|
| GO:0048870~cell motility | 52 | HRAS, PDGFA, MMP9, FERMT3, FERMT1, SDC4, CD151, AMOTL1, WNT4, APOE, SEMA3F, ADAM8, NRG1, FAM83H, DAB2IP, HYAL2, FOXJ1, PDPN, PLXNB2, EFNB1, MYH9, JUP, TACSTD2, VEGFA, CEMIP, SEMA4C, SEMA4B, MDM2, LAMC2, WNT11, MYO18A, ALOX12, SORD, DAG1, ITGB4, CX3CL1, CALR, SRC, P2RY1, FAT2, CNN2, HSPA5, LAMB1, PLTP, COL18A1, MYO1C, TGFBR1, ITGA3, EPHA1, GAS6, ITGA6, LAMA5 | 3,28 | 5,74E-11 |
| GO:0016477~cell migration | 50 | HRAS, PDGFA, MMP9, FERMT3, FERMT1, SDC4, CD151, AMOTL1, WNT4, APOE, SEMA3F, NRG1, FAM83H, ADAM8, DAB2IP, HYAL2, FOXJ1, PDPN, PLXNB2, EFNB1, MYH9, JUP, TACSTD2, VEGFA, SEMA4C, CEMIP, SEMA4B, MDM2, LAMC2, WNT11, MYO18A, ALOX12, DAG1, ITGB4, CX3CL1, CALR, SRC, P2RY1, FAT2, CNN2, HSPA5, LAMB1, COL18A1, MYO1C, TGFBR1, ITGA3, EPHA1, GAS6, ITGA6, LAMA5 | 3,54 | 1,08E-11 |
| GO:2000026~regulation of multicellular organismal development | 50 | PDGFA, MMP9, WNT3A, ITPKB, WNT4, GATA6, APOE, SEMA3F, EEF2K, NRG1, ADAM8, DAB2IP, SLC25A4, FOXJ1, PLXNB2, KRT17, TACSTD2, VEGFA, SEMA4C, SEMA4B, MDM2, WNT11, WNT9A, NGFR, TRP73, ALOX12, PHLDB1, DAG1, CD109, SFN, CX3CL1, CALR, ZFP36L2, CYP26B1, BCL9L, HSPA5, MAFB, IL1RL2, TGFBR1, CD276, ITGA3, EPHA1, GAS6, RGS14, CDKN1C, WNT7B, LAMA5, ID1, BMP7, BMP6 | 2,25 | 9,77E-05 |
| GO:0050790~regulation of catalytic activity | 48 | FXYD3, HRAS, PDGFA, HIP1R, WNT3A, MMP9, CD109, DAG1, CX3CL1, SFN, FEM1B, SDC4, 1300017J02RIK, SRC, PXN, WNT4, COL7A1, APOE, TGFA, HSPA5, ADAM8, NRG1, MAPKBP1, DAB2IP, HYAL2, FOXJ1, PLXNB2, TGFBR1, RCAN1, EPHA1, RGS14, GAS6, CARD10, WNT7B, HSP90B1, ITGA6, VEGFA, CEMIP, SETD7, MDM2, WNT11, TRP73, NGFR, WNT9A, BMP7, PERP, DUSP7, ALOX12 | 2,45 | 1,41E-05 |
| GO:0098609~cell-cell adhesion | 46 | LIMA1, ASS1, FERMT3, WNT3A, JAG2, PDIA6, ITPKB, CX3CL1, SFN, TAGLN2, SDC4, CD151, SRC, PAK6, ZFP36L2, WNT4, FAT2, CYP26B1, IDH1, CNN2, FAM129B, ADAM8, LAMB1, ERF, FOXJ1, PDPN, MAFB, EFNB1, PLXNB2, IL1RL2, CD276, CSRP1, MYH9, FLNB, PPP1R13L, JUP, WNT7B, ITGA6, TACSTD2, PKP3, CAPG, VEGFA, DSP, BMP7, PERP, ALOX12 | 3,55 | 1,74E-10 |
| GO:0051270~regulation of cellular component movement | 44 | HRAS, PDGFA, FERMT3, MMP9, WNT3A, DAG1, CX3CL1, CALR, AMOTL1, SDC4, SRC, WNT4, APOE, SEMA3F, CNN2, HSPA5, ADAM8, FAM83H, NRG1, LAMB1, COL18A1, DAB2IP, HYAL2, MYO1C, PDPN, PLXNB2, TGFBR1, CHST3, ITGA3, EPHA1, GAS6, JUP, ITGA6, TACSTD2, LAMA5, VEGFA, CEMIP, SEMA4C, DSP, MDM2, SEMA4B, LAMC2, WNT11, ALOX12 | 4,42 | 3,44E-13 |

| | | | | |
|---|---|---|---|---|
| GO:0030030~cell projection organization | 41 | LIMA1, HRAS, PDGFA, WNT3A, DAG1, ITGB4, SDC4, SRC, PXN, APOE, SEMA3F, POU4F3, EEF2K, HSPA5, NRG1, LAMB1, DAB2IP, FOXJ1, EFNB1, PLXNB2, TGFBR1, ITGA3, MYH9, MARK4, MICALL1, WNT7B, NPTXR, ITGA6, TACSTD2, ID1, LAMA5, VEGFA, CAPG, SEMA4C, MDM2, SEMA4B, NGFR, BMP7, DST, EPS8L2, SEPT9 | 2,50 | 1,50E-04 |
| GO:2000145~regulation of cell motility | 40 | HRAS, PDGFA, FERMT3, MMP9, DAG1, CX3CL1, CALR, AMOTL1, SDC4, SRC, WNT4, APOE, SEMA3F, CNN2, HSPA5, ADAM8, FAM83H, NRG1, LAMB1, COL18A1, DAB2IP, HYAL2, MYO1C, PDPN, PLXNB2, TGFBR1, ITGA3, EPHA1, GAS6, ITGA6, LAMA5, TACSTD2, VEGFA, CEMIP, SEMA4C, MDM2, SEMA4B, LAMC2, WNT11, ALOX12 | 4,34 | 1,96E-11 |
| GO:0098602~single organism cell adhesion | 37 | ASS1, FERMT3, WNT3A, JAG2, PDIA6, ITPKB, CALR, SDC4, CD151, PXN, SRC, ZFP36L2, WNT4, CYP26B1, ADAM8, LAMB1, ERF, FOXJ1, PDPN, MAFB, EFNB1, IL1RL2, CD276, CSRP1, MYH9, EPHA1, JUP, WNT7B, ITGA6, LAMA5, TACSTD2, PKP3, VEGFA, DSP, BMP7, PERP, ALOX12 | 3,95 | 4,19E-09 |
| GO:0009968~negative regulation of signal transduction | 33 | MMP9, WNT3A, DAG1, CD109, CX3CL1, CALR, SRC, WNT4, APOE, CYP26B1, BCL9L, HSPA5, DAB2IP, HYAL2, TGFBR1, ACKR3, ITGA3, RGS14, GAS6, HYOU1, DHRS3, WNT7B, NXN, ITGA6, RGS3, VEGFA, MDM2, WNT11, TRP73, NGFR, WNT9A, BMP7, DUSP7 | 2,73 | 5,82E-04 |
| GO:0010648~negative regulation of cell communication | 33 | MMP9, WNT3A, DAG1, CD109, CX3CL1, CALR, SRC, WNT4, APOE, CYP26B1, BCL9L, HSPA5, DAB2IP, HYAL2, TGFBR1, ACKR3, ITGA3, RGS14, GAS6, HYOU1, DHRS3, WNT7B, NXN, ITGA6, RGS3, VEGFA, MDM2, WNT11, TRP73, NGFR, WNT9A, BMP7, DUSP7 | 2,42 | 8,22E-03 |
| GO:0023057~negative regulation of signaling | 33 | MMP9, WNT3A, DAG1, CD109, CX3CL1, CALR, SRC, WNT4, APOE, CYP26B1, BCL9L, HSPA5, DAB2IP, HYAL2, TGFBR1, ACKR3, ITGA3, RGS14, GAS6, HYOU1, DHRS3, WNT7B, NXN, ITGA6, RGS3, VEGFA, MDM2, WNT11, TRP73, NGFR, WNT9A, BMP7, DUSP7 | 2,41 | 8,82E-03 |
| GO:0016337~single organismal cell-cell adhesion | 32 | ASS1, FERMT3, WNT3A, JAG2, PDIA6, ITPKB, SDC4, CD151, SRC, ZFP36L2, WNT4, CYP26B1, ADAM8, LAMB1, ERF, FOXJ1, PDPN, MAFB, EFNB1, IL1RL2, CD276, CSRP1, MYH9, JUP, WNT7B, ITGA6, PKP3, VEGFA, DSP, PERP, BMP7, ALOX12 | 3,68 | 1,02E-06 |
| GO:0051272~positive regulation of cellular component movement | 31 | HRAS, PDGFA, FERMT3, MMP9, CX3CL1, CALR, AMOTL1, SRC, SEMA3F, HSPA5, ADAM8, FAM83H, LAMB1, COL18A1, DAB2IP, MYO1C, PDPN, TGFBR1, CHST3, ITGA3, EPHA1, GAS6, ITGA6, VEGFA, CEMIP, SEMA4C, SEMA4B, MDM2, LAMC2, WNT11, ALOX12 | 5,58 | 6,51E-11 |

| | | | | |
|---|---|---|---|---|
| GO:2000147~positive regulation of cell motility | 30 | HRAS, PDGFA, FERMT3, MMP9, CX3CL1, CALR, AMOTL1, SRC, SEMA3F, HSPA5, ADAM8, FAM83H, LAMB1, COL18A1, DAB2IP, MYO1C, PDPN, TGFBR1, ITGA3, EPHA1, GAS6, ITGA6, VEGFA, CEMIP, SEMA4C, SEMA4B, MDM2, LAMC2, WNT11, ALOX12 | 5,53 | 2,34E-10 |
| GO:0040017~positive regulation of locomotion | 30 | HRAS, PDGFA, FERMT3, MMP9, CX3CL1, CALR, AMOTL1, SRC, SEMA3F, HSPA5, ADAM8, FAM83H, LAMB1, COL18A1, DAB2IP, MYO1C, PDPN, TGFBR1, ITGA3, EPHA1, GAS6, ITGA6, VEGFA, CEMIP, SEMA4C, SEMA4B, MDM2, LAMC2, WNT11, ALOX12 | 5,37 | 4,71E-10 |
| GO:0030029~actin filament-based process | 28 | HRAS, LIMA1, PDGFA, HIP1R, CALR, SDC4, SRC, WNT4, CNN2, MYO1C, FOXJ1, TGFBR1, CORO6, CORO7, CSRP1, MYH9, EPHA1, FLNB, JUP, HSP90B1, KRT19, TACSTD2, ID1, CAPG, SCN4B, DSP, WNT11, MYO18A | 3,62 | 2,21E-05 |
| GO:0001568~blood vessel development | 27 | PDGFA, MMP9, CX3CL1, WNT4, GATA6, APOE, TGFA, HS6ST1, ADAM8, PCSK5, COL18A1, DAB2IP, TGFBR1, HSPG2, ACKR3, MFGE8, MYH9, EPHA1, JUNB, DLX3, WNT7B, VEGFA, MDM2, WNT11, NGFR, BMP7, ALOX12 | 3,58 | 5,62E-05 |
| GO:0048514~blood vessel morphogenesis | 23 | COL18A1, DAB2IP, PDGFA, TGFBR1, MMP9, HSPG2, ACKR3, MFGE8, CX3CL1, MYH9, EPHA1, JUNB, WNT4, WNT7B, GATA6, APOE, VEGFA, TGFA, WNT11, HS6ST1, NGFR, ADAM8, ALOX12 | 3,62 | 6,32E-04 |
| GO:0031589~cell-substrate adhesion | 21 | FERMT3, ITGB4, DAG1, BCAM, ITGA3, NID2, CALR, SDC4, EPHA1, SRC, PXN, GAS6, WNT4, ITGA6, TACSTD2, LAMA5, ID1, VEGFA, ADAM8, LAMB1, VWA2 | 6,00 | 5,32E-07 |
| GO:0001525~angiogenesis | 19 | COL18A1, DAB2IP, PDGFA, TGFBR1, MMP9, HSPG2, ACKR3, MFGE8, CX3CL1, MYH9, EPHA1, WNT7B, GATA6, VEGFA, TGFA, HS6ST1, NGFR, ADAM8, ALOX12 | 3,63 | 8,08E-03 |
| GO:0045785~positive regulation of cell adhesion | 18 | WNT3A, IL1RL2, EFNB1, DAG1, CD276, ITPKB, ITGA3, CX3CL1, CALR, SDC4, EPHA1, SRC, WNT4, ITGA6, VEGFA, ADAM8, BMP7, ALOX12 | 4,18 | 2,43E-03 |
| GO:0022407~regulation of cell-cell adhesion | 17 | ASS1, FOXJ1, FERMT3, WNT3A, EFNB1, IL1RL2, CD276, ITPKB, SDC4, SRC, WNT4, ITGA6, VEGFA, CYP26B1, ADAM8, BMP7, ALOX12 | 3,80 | 1,72E-02 |
| GO:0032970~regulation of actin filament-based process | 16 | LIMA1, HRAS, MYO1C, HIP1R, TGFBR1, MYH9, SDC4, EPHA1, JUP, WNT4, TACSTD2, ID1, CAPG, DSP, WNT11, CNN2 | 3,94 | 2,23E-02 |
| GO:0034330~cell junction organization | 14 | MYO1C, TGFBR1, ITGB4, PTPN13, SDC4, SRC, PXN, JUP, WNT4, PKP3, VEGFA, DSP, WNT11, PERP | 5,97 | 1,03E-03 |
| GO:0043062~extracellular structure organization | 14 | COL18A1, PHLDB1, MMP9, WNT3A, TGFBR1, DAG1, HSPG2, NID2, GAS6, LAMB3, LAMA5, LAMC2, VWA1, LAMB1 | 5,19 | 4,95E-03 |
